# Identification of Potential Therapeutic Targets and Biomarkers for Glioblastomas Through Integrative Analysis of Gene Expression Data

**DOI:** 10.1101/2024.03.04.583392

**Authors:** Angélica Bautista, Ricardo Romero

## Abstract

**Background:** In this study, we conducted a comprehensive analysis of differential gene expression data from studies GSE15824, GSE4290 and GEPIA2 data to identify up-regulated hub genes with potential as therapeutic targets for glioblastomas. Through virtual screening, we also aimed to identify novel VEGFA inhibitors.

**Results:** Seven up-regulated hub genes (TYROBP, ITGB2, C1QA, C1QB, CTSS, TLR2, and CD163) were identified. Virtual screening of VEGFA inhibitors led to the discovery of six significant hits, including three from the ChemDiv library (D519-0372, G868-0191, and Y031-5201) and three from the ZINC20 database (ZINC57658, ZINC57652, ZINC57679). Molecular dynamics simulations highlighted G868-0191 as the most stable VEGFA inhibitor. Two repurposed drugs, Sunitinib and Ticlopidine hydrochloride, were also identified as potential candidates. In addition, the down-regulated hub genes GABARAPL1, OPTN, and CDH8 were proposed as potential biomarkers for glioblastomas.

**Conclusion:** This study underscores the significance of immune-related hub genes in glioblastoma pathology and suggests new VEGFA inhibitors as promising therapeutic agents. The identification of down-regulated genes as potential biomarkers offers further avenues for clinical application. However, experimental validation is needed to confirm the clinical utility of these findings.

## Introduction

Malignant tumors of the central nervous system (CNS) are characterized by their aggressive nature and poor prognosis. Among these tumors, gliomas are the most common primary CNS tumors, exhibiting the ability to differentiate into various cell types, such as neural stem cells, astrocytes, or oligodendroglia progenitor cells [1].

Glioblastomas, classified as level IV gliomas by the World Health Organization (WHO), are the most aggressive subtype, accounting for 60-70% of malignant gliomas [2,3]. They can be further categorized into three subtypes: proneural, classic, and mesenchymal [4]. Globally, the incidence of glioblastoma is approximately 3.19 per 100,000 person-years, with higher rates observed in developed countries [5]. The incidence of gliomas increases with age, particularly among individuals over 75 years old, with age also correlating inversely with survival rate [6]. While the exact risk factors for glioblastoma development remain unclear, it has been observed that it is more common in men than in women, with a male-to-female ratio of 1.6:1 [7–9].

Known risk factors for glioblastoma include exposure to high-dose ionizing radiation and certain genetic conditions such as neurofibromatosis type 1, tuberous sclerosis, and Li-Fraumeni syndrome [10]. Other potential risk factors under investigation include occupational exposures, electromagnetic fields, and certain viruses, although their roles remain controversial [11].

Vascular Endothelial Growth Factor A (VEGFA) has been identified as a validated target for glioblastoma treatment due to its crucial role in tumor angiogenesis [12–19]. However, current VEGFA inhibitors, such as bevacizumab, have shown limited efficacy on overall survival and are associated with significant side effects and the development of resistance mechanisms [20–23]. Therefore, there is an urgent need for alternative inhibitors with improved efficacy and safety profiles.

In silico analyses enable the identification of compounds that may possess inhibitory effects on overexpressed proteins associated with various diseases and malignancies, such as glioblastomas. They allow for the evaluation of the stability of interactions between candidate compounds and proteins, and offer cost-effective alternatives by studying existing drugs, as compared to the expenses involved in discovering new compounds. Moreover, with the availability of databases containing drugs with inhibitory potential against numerous genes, as well as the integration of Artificial Intelligence, the search for candidate compounds can be expedited [24–27].

The purpose of the present work is two-fold:

1. To provide alternative inhibitors of the validated glioblastoma target VEGFA, highlighting the shortcomings of current inhibitors. This will be achieved through virtual screening, involving both molecular docking and ADMET filtering, of small molecule databases.
2. To identify other potential therapeutic targets through gene expression and integrative bioinformatic analyses from two databases: The Cancer Genome Atlas (TCGA) [28] and the Gene Expression Omnibus (GEO). This approach will allow for a comprehensive exploration of novel targets and potential combination therapies.

The study will also include screening of control drugs for identified targets and enrichment analysis of hub genes to complement the findings. This two-pronged approach—targeting VEGFA and discovering new targets—aims to expand the therapeutic landscape for glioblastomas and address the urgent need for more effective treatments.

## Methods and Materials

### Differential gene expression analyses

To identify hub genes associated with glioblastomas, gene expression analyses were performed on samples from two databases, Gene Expression Omnibus (GEO) and GEPIA2. From the former two studies were selected. One study, “Gene expression profiling of human gliomas and human glioblastoma cell lines,” [29] with accession number GSE15824, utilized microarray analysis on 30 brain tumor samples (12 primary glioblastomas, 3 secondary glioblastomas, 8 astrocytoma, and 7 oligodendrogliomas) and 5 cell lines (LN018, LN215, LN229, LN319, and BS149) to identify differentially expressed signaling pathways in human gliomas compared to normal brain tissue and normal astrocytes.

The other study, “Expression data of glioma samples from Henry Ford Hospital,” [30] with accession number GSE4290, evaluated mRNA expression of 153 samples from patients with brain tumors (77 glioblastomas, 50 oligodendrogliomas, and 26 astrocytoma), with 23 samples from patients with epilepsy without tumors serving as controls.

The analyses were performed using NCBI’s online tool GEO2R, with options Benjamini & Hochberg for false discovery rate, log transformation, limma precision weights and forced normalization. Adjusted P value significance cut-off was set to 0.05 and log2 fold change threshold was set to zero. Common up and down-regulated genes from both studies were identified using the Funrich [31–33] software.

In parallel, differential gene expression analysis was performed on tumor vs normal glioblastoma multiforme (GBM) samples from GEPIA2 [34], which integrates data from The Cancer Genome Atlas Program (TCGA) and The Genotype-Tissue Expression (GTEx) project [35]. The analysis was conducted on the GEPIA2 portal, with a |Log2FC| cutoff of 1 and a q-value (adjusted p value) cutoff of 0.05, with the LIMMA method.

### Analysis of Interaction Network

The up and down-regulated genes common to both databases (GEPIA2 and the common genes from both GEO studies) were identified using Funrich, and the protein – protein interaction (PPI) network among the proteins codified by those genes was obtained using the STRING database (Ver. 12) [36, 37] with a high confidence interaction score of 0.7 for the upregulated genes. The top dysregulated hub genes, chosen by both degree of interaction and betweenness centrality metrics, were subsequently identified using the Cytohubba [38] plugin in the Cytoscape software [39]. Survival analyses were also performed for hub genes against glioblastoma multiforme (GBM) and low-grade gliomas (LGG) data from GEPIA2.

### Preparation of Receptor

The validated target Vascular Endothelial Growth Factor A (VEGFA) protein was used for virtual screening of alternative drug candidates. The human protein sequence was searched for in the Uniprot database, and a BLAST search was performed using the VEGFA sequence against the Protein Data Bank. The protein with entry 1VPF [40] (Figure1), which had a percentage of identity greater than 80% and an unmutated structure, was selected. Cavity detection was conducted in CASTp [41] and the structure was prepared for molecular docking by removing non-protein chains, solvents, ions, and ligands in Chimera [42]. Energy minimization was performed in Swiss PDB viewer [43], which includes a partial implementation of the GROMOS96 force-field, and the minimization is conducted in vacuo by the steepest descent method [44]. Addition of hydrogens and charges to the protein structure were performed in Chimera.

**Figure 1.**
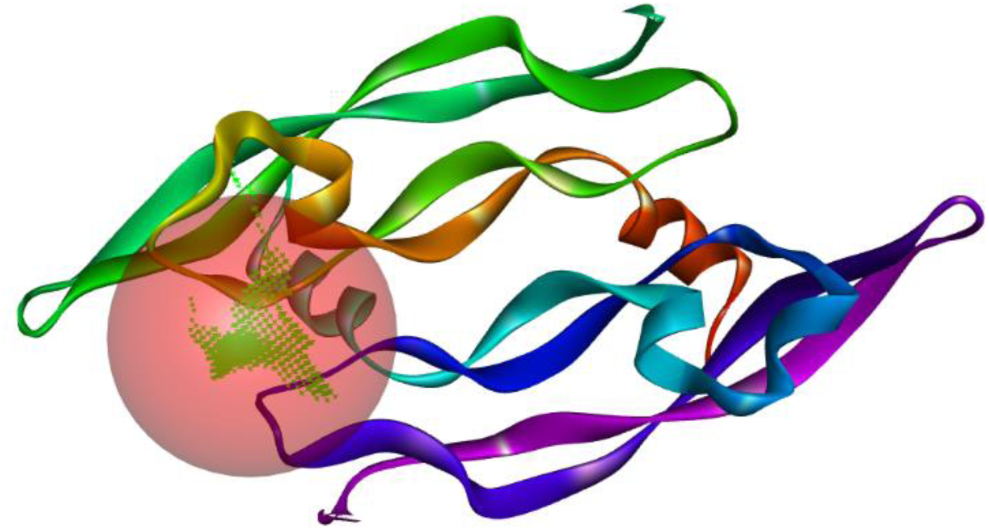
VEGFA Structure PDB 1VPF (chains A and B), showing the pocket for the active site, encompassing both chains.

### Virtual screening

Ligands were obtained from the ZINC20 database, 100 compounds were downloaded from the subset of inferred compounds targeting the protein VEGFA, and from the Protein Kinases Inhibitors Library from ChemDiv [45], which consists of 36324 compounds. Energy minimization was performed with the Open Babel software [46], using the generalized Amber force field (GAFF) with 5000 steps by steepest descent with linear convergence.

The Drug Gene Interaction Database (https://www.dgidb.org/), with VEGFA as target, was also searched for drugs repurposing. Approved drugs Irinotecan, Carboplatin and Fluorouracil were used as controls, based on information about their use in glioblastoma treatment and VEGFA interaction, retrieved from the Drug Bank (https://go.drugbank.com/) and Gene Cards (https://www.genecards.org) databases.

Molecular docking was conducted with Autodock Vina [47, 48], with center coordinates (37, 11, 14) and a symmetric grid size of 25 Å, and the results were filtered for druglikeness and toxicity properties using SwissADME [49], OSIRIS Property Explorer and Data Warrior [50]. Finalists’ compounds were required to pass all the druglikeness rules of Lipinski [51], Ghose [52], Veber [53], Egan [54], Muegge [55], exhibit none of the structure Brenk [56] and PAINS alerts [57] and permeate the Brain Blood Barrier (BBB). As for toxicity, they were required to show zero alerts for mutagenicity, tumorigenicity, irritancy, and reproductive effects. They were also filtered by their bioactivity, using a deep learning QSAR model trained on tyrosine kinase inhibitors. [58,59]

Using values provided by the OSIRIS Property Explorer for druglikeness, drug-score, and solubility, as well as the binding energy, in absolute values, obtained from docking, a simple average score was calculated to rank the final hits.

### Molecular Dynamics Simulations

Molecular dynamics simulations were conducted for the best scoring compounds from the ChemDiv library. They were run for 30 ns using GROMACS,[60] with the CHARMM27 force field and TIP3P solvation. Given that the protein’s active site involves both chains, we used normal mode analysis to obtain distance restraints to stabilize the complexes.

### Enrichment Analysis

Enrichment analysis was performed on the hub genes using GeneTrail [61] to obtain the ten most enriched categories according to the KEGG [62], Wikipathways [63], Reactome [64] and Gene Ontology [65, 66] databases.

## Results

We obtained six significant hits from virtual screening, three from the ChemDiv library with identifiers D519-0372, G868-0191, and Y031-5201, which presented the highest scores and best overall properties, and three from the ZINC20 database, with identifiers ZINC57658, ZINC57652 and ZINC57679. These hits outperformed control drugs in binding energy, toxicity and druglikeness properties, and among them the three compounds from the ChemDiv library attained the highest scores, so they were selected for further investigation with molecular dynamics. We also identified two drugs for repurposing: Sunitinib and Ticlopidine hydrochloride. Compounds’ properties are provided in Tables I and II, as well as their druglikeness and toxicity results, along with the controls.

**Table I.** SMILES strings, molecular weight (MW) in g/mol, topological surface area (TPSA) in Å^2^, and brain blood barrier permeability (BBB) for the finalists’ hits, control and repurpose drugs. The Ligand column shows the name provided by PubChem.

| Id | Ligand | SMILES | MW | TPSA | BBB |
| --- | --- | --- | --- | --- | --- |
| Sunitinib | Sunitinib | CCN(CCNC(=O)c1c(C)[nH]c(c1C)C=C1C(=O)Nc2c1cc(F)cc2)CC | 398.47 | 77.23 | Yes |
| D519-0372 | AKOS004929708 | CN1c(cc(cc2)NC(c3noc(-c4ccccc4)c3)=O)c2OCC1=O | 349.35 | 84.67 | Yes |
| G868-0191 | - | CC(N(CC1)CCN1c1nc(Nc2ccc(C)cc2)cc(C)n1)=O | 325.40 | 61.36 | Yes |
| Y031-5201 | - | Cc1cc(NC(C(CC2=O)CN2c2cc(C)cc(C)c2)=O)ncc1 | 323.40 | 62.30 | Yes |
| Ticlopidine | Ticlopidine hydrochloride | Clc1ccccc1CN1CCc2c(C1)ccs2.Cl | 300.25 | 31.48 | Yes |
| ZINC57679 | 4'-Hydroxyflavanone | Oc1ccc(cc1)c1cc(=O)c2c(o1)cccc2 | 238.24 | 50.44 | Yes |
| ZINC57652 | 5,4'-Dihydroxyflavone | Oc1ccc(cc1)c1cc(=O)c2c(o1)cccc2O | 254.24 | 70.67 | Yes |
| Control | Irinotecan | CCC1(O)C(=O)OCc2c1cc1c3nc4ccc(cc4c(c3Cn1c2=O)CC)OC(=O)N1CCC(CC1)N1CCCCC1 | 586.68 | 114.2 | No |
| ZINC57658 | Liquiritigenin | Oc1ccc(cc1)c1cc(=O)c2c(o1)cc(cc2)O | 254.24 | 70.67 | Yes |
| Control | Carboplatin | OC(=O)C1(CCC1)C(=O)O.[Pt+2].[NH2-].[NH2-] | 371.25 | 126.64 | No |
| Control | Fluorouracil | Fc1c[nH]c(=O)[nH]c1=O | 130.08 | 65.72 | No |

**Table II.** Druglikenes and toxicity properties. The columns Lipinski (L), Ghose (G), Veber (V), Egan (E) and Muegge (M) show the number of violations to the corresponding rules. The columns PAINS and Brenk show the number of alerts. Toxicity includes information about mutagenicity, tumorigenic, irritant, and reproductive effectiveness, provided by Osiris Property Explorer, while Solubility, Druglikeness, and Drug score are the scores provided by the same software. Energy is the binding energy reported by docking, in kcal/mol, and the final Score is the simple arithmetic mean of the preceding four columns, with Energy in absolute value. All the compounds were classified as active by the deep neural network model.

| Ligand | L | G | V | E | M | PAINS | Brenk | Toxicity | Energy | Solubility | Druglikeness | Drug score | Score |
| --- | --- | --- | --- | --- | --- | --- | --- | --- | --- | --- | --- | --- | --- |
| Sunitinib | 0 | 0 | 0 | 0 | 0 | 0 | 1 | Irritant | -7.07 | -3.47 | 8.47 | 0.47 | 3.13 |
| D519-0372 | 0 | 0 | 0 | 0 | 0 | 0 | 0 | None | -8.71 | -4.21 | 5.92 | 0.76 | 2.79 |
| G868-0191 | 0 | 0 | 0 | 0 | 0 | 0 | 0 | None | -8.43 | -3.58 | 4.82 | 0.8 | 2.62 |
| Y031-5201 | 0 | 0 | 0 | 0 | 0 | 0 | 0 | None | -8.17 | -4.22 | 1.22 | 0.68 | 1.46 |
| Ticlopidine hydrochloride | 0 | 0 | 0 | 0 | 0 | 0 | 0 | None | -6.08 | -3.55 | 4.68 | 0.81 | 2.01 |
| ZINC57679 | 0 | 0 | 0 | 0 | 0 | 0 | 0 | None | -7.48 | -3.45 | 2.11 | 0.79 | 1.73 |
| ZINC57652 | 0 | 0 | 0 | 0 | 0 | 0 | 0 | None | -7.66 | -3.15 | 1.56 | 0.79 | 1.71 |
| Irinotecan | 1 | 3 | 0 | 0 | 0 | 0 | 0 | None | -8.98 | -4.5 | 0.07 | 0.35 | 1.22 |
| ZINC57658 | 0 | 0 | 0 | 0 | 0 | 0 | 0 | None | -7.75 | -3.15 | -0.36 | 0.61 | 1.21 |
| Carboplatin | 0 | 0 | 0 | 0 | 1 | 0 | 1 | None | -4.93 | -0.9 | -5.33 | 0.49 | -0.2 |
| Fluorouracil | 0 | 3 | 0 | 0 | 2 | 0 | 0 | All | -4.46 | -1.76 | -4.5 | 0.06 | -0.43 |

The Venn diagrams of the dysregulated genes from both databases are presented in Figure 2, showing they share 185 genes for the former case and 151 for the latter. From these, the PPI networks of the associated proteins were obtained from Strings DB (Figures in supplementary material), with significant enrichment p values, showing that the networks constitute biological clusters. Finally, dysregulated hub genes are given in Figure 3.

**Figure 2.**
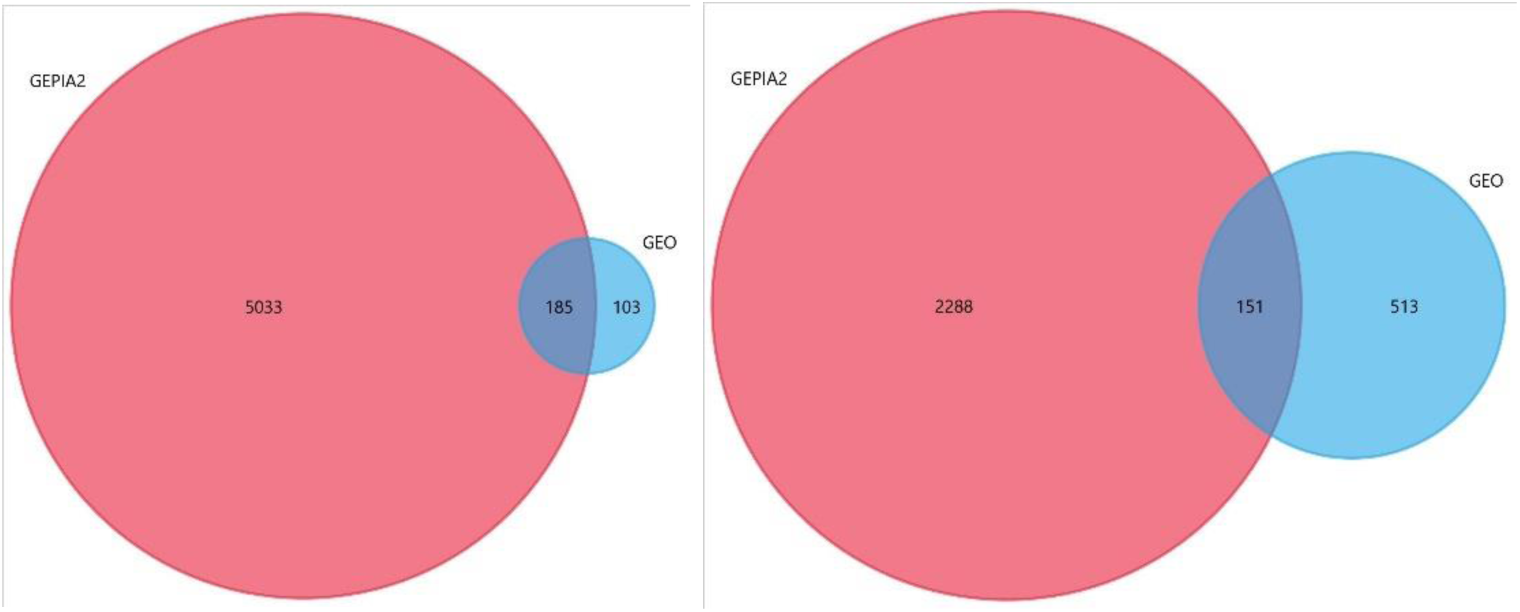
Venn Diagrams showing the number of common genes from GEO studies and GEPIA2 for up-regulated genes (left) and down-regulated ones (right).

**Figure 3.**
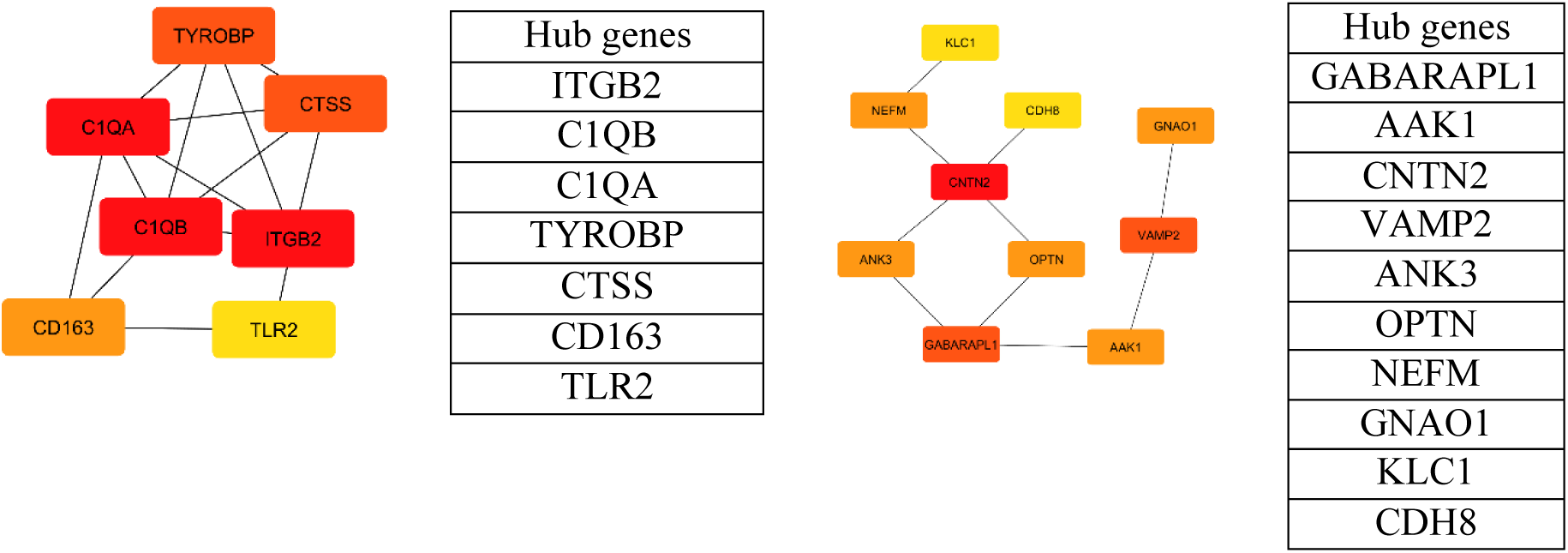
Up-regulated (left) and down regulated (right) hub genes obtained from the PPI networks of common genes from GEO and GEPIA. Both degree and betweenness centrality metrics were used to determine them. Colors represent the interaction degree from red (highest) to yellow (lowest).

Docking results from the ChemDiv and ZINC compounds and controls are presented in Tables III and IV, and the docked complexes for the former in Figures 4 to 6. Figures 7-12 present the results from the molecular dynamics simulations of the three hits from ChemDiv in complex with VEGFA, including RMSD, RMSF, potential energy and hydrogen bonds plots. Results from the enrichment and survival analyses are given in Figures 13 and 14, respectively.

**Table III.** Docked complexes interactions between hits from ChemDiv and VEGFA (1VPF)

| Compound | Binding energy (Kcal/mol) | Donnor | Distance (Å) | Acceptor | Category | Interaction type |
| --- | --- | --- | --- | --- | --- | --- |
| D519-0372 |  | :UNL1:C | 3.712940 | B:ASP34:OD2 | Hydrogen Bond | Carbon Hydrogen Bond |
|  |  | :UNL1 | 4.861817 | A:CYS60 | Hydrophobic | Pi-Alkyl |
|  |  | A:CYS61:N | 4.033804 | :UNL1 | Hydrogen Bond | Pi-Donor Hydrogen Bond |
|  |  | :UNL1 | 5.026478 | A:CYS61 | Hydrophobic | Pi-Alkyl |
|  |  | :UNL1:H | 2.521557 | A:CYS61:O | Hydrogen Bond | Conventional Hydrogen Bond |
|  |  | :UNL1 | 5.343835 | B:ILE46 | Hydrophobic | Pi-Alkyl |
|  |  | :UNL1:H | 2.189874 | B:SER50:OG | Hydrogen Bond | Conventional Hydrogen Bond |
|  |  | A:ASP63:C,O; GLU64:N | 3.726017 | :UNL1 | Hydrophobic | Amide-Pi Stacked |
|  |  | :UNL1 | 4.699170 | B:ILE46 | Hydrophobic | Pi-Alkyl |
|  |  | A:ASN62:C,O; ASP63:N | 3.883579 | :UNL1 | Hydrophobic | Amide-Pi Stacked |
|  |  | :UNL1 | 5.187501 | B:LYS48 | Hydrophobic | Pi-Alkyl |
| G868-0191 |  | A:LYS107:NZ | 3.008114 | :UNL1:O | Hydrogen Bond | Conventional Hydrogen Bond |
|  |  | :UNL1:C | 3.449608 | A:GLU64:OE2 | Hydrogen Bond | Carbon Hydrogen Bond |
|  |  | :UNL1:C | 3.243453 | A:ASP63:O | Hydrogen Bond | Carbon Hydrogen Bond |
|  |  | :UNL1:C | 2.920549 | A:CYS61:O | Hydrogen Bond | Carbon Hydrogen Bond |
|  |  | A:ASN62:C,O; ASP63:N | 3.535251 | :UNL1 | Hydrophobic | Amide-Pi Stacked |
|  |  | :UNL1 | 4.956026 | B:LYS48 | Hydrophobic | Pi-Alkyl |
|  |  | :UNL1 | 5.226545 | B:ILE46 | Hydrophobic | Pi-Alkyl |
|  |  | :UNL1 | 5.176662 | B:ILE46 | Hydrophobic | Pi-Alkyl |
|  |  | :UNL1:C | 4.219850 | B:ILE46 | Hydrophobic | Alkyl |
|  |  | B:PHE36 | 4.981074 | :UNL1:C | Hydrophobic | Pi-Alkyl |
|  |  | :UNL1:H | 2.431903 | B:PHE47:O | Hydrogen Bond | Conventional Hydrogen Bond |
| Y031-5201 |  | :UNL1:C | 3.689966 | B:LYS48 | Hydrophobic | Alkyl |
|  |  | A:ASN62:C,O; ASP63:N | 4.022892 | :UNL1 | Hydrophobic | Amide-Pi Stacked |
|  |  | B:ILE46:CG2 | 3.496250 | :UNL1 | Hydrophobic | Pi-Sigma |
|  |  | :UNL1:C | 4.548717 | B:ILE46 | Hydrophobic | Alkyl |
|  |  | B:SER50:OG | 2.975823 | :UNL1:O | Hydrogen Bond | Conventional Hydrogen Bond |

**Table IV.** Docked complexes interactions between flavonoids from ZINC and VEGFA (1VPF)

| Compound | Binding energy (Kcal/mol) | Donnor | Distance (Å) | Acceptor | Category | Interaction type |
| --- | --- | --- | --- | --- | --- | --- |
| ZINC57652 | -7.66 | A:GLU64:OE2 | 4.57 | UNL1 | Electrostatic | Pi-Anion |
|  |  | A:GLU64:CA | 4 | UNL1 | Hydrophobic | Pi-Sigma |
|  |  | A:ASP63:C,O;GLU64:N | 4.36 | UNL1 | Hydrophobic | Amide-Pi Stacked |
|  |  | A:ASP63:C,O;GLU64:N | 4.05 | UNL1 | Hydrophobic | Amide-Pi Stacked |
|  |  | A:ASP63:N | 3.34 | UNL1:O | Hydrogen Bond | Conventional Hydrogen Bond |
|  |  | A:ASN62:C,O;ASP63:N | 3.6 | UNL1 | Hydrophobic | Amide-Pi Stacked |
|  |  | UNL1 | 5.13 | B:ILE46 | Hydrophobic | Pi-Alkyl |
|  |  | UNL1 | 5.03 | B:ILE46 | Hydrophobic | Pi-Alkyl |
|  |  | UNL1 | 5 | B:LYS48 | Hydrophobic | Pi-Alkyl |
|  |  | B:LYS48:NZ | 3.17 | UNL1:O | Hydrogen Bond | Conventional Hydrogen Bond |
| ZINC57658 | -7.75 | A:GLU64:OE2 | 4.63 | UNL1 | Electrostatic | Pi-Anion |
|  |  | A:ASP63:C,O;GLU64:N | 4.36 | UNL1 | Hydrophobic | Amide-Pi Stacked |
|  |  | A:ASP63:C,O;GLU64:N | 4.04 | UNL1 | Hydrophobic | Amide-Pi Stacked |
|  |  | A:ASP63:N | 3.32 | UNL1:O | Hydrogen Bond | Conventional Hydrogen Bond |
|  |  | A:ASN62:C,O;ASP63:N | 3.6 | UNL1 | Hydrophobic | Amide-Pi Stacked |
|  |  | B:LYS48:NZ | 3.27 | UNL1:O | Hydrogen Bond | Conventional Hydrogen Bond |
|  |  | UNL1 | 5.11 | B:ILE46 | Hydrophobic | Pi-Alkyl |
|  |  | UNL1 | 5.02 | B:ILE46 | Hydrophobic | Pi-Alkyl |
|  |  | UNL1 | 5 | B:LYS48 | Hydrophobic | Pi-Alkyl |
| ZINC57679 | -7.48 | UNL1 | 5.1 | B:ILE46 | Hydrophobic | Pi-Alkyl |
|  |  | UNL1 | 5.06 | B:ILE46 | Hydrophobic | Pi-Alkyl |
|  |  | UNL1 | 5 | B:LYS48 | Hydrophobic | Pi-Alkyl |
|  |  | A:GLU64:OE2 | 4.6 | UNL1 | Electrostatic | Pi-Anion |
|  |  | A:GLU64:CA | 4 | UNL1 | Hydrophobic | Pi-Sigma |
|  |  | A:ASP63:C,O;GLU64:N | 4.35 | UNL1 | Hydrophobic | Amide-Pi Stacked |
|  |  | A:ASP63:C,O;GLU64:N | 4.05 | UNL1 | Hydrophobic | Amide-Pi Stacked |
|  |  | A:ASP63:N | 3.32 | UNL1:O | Hydrogen Bond | Conventional Hydrogen Bond |
|  |  | A:ASN62:C,O;ASP63:N | 3.6 | UNL1 | Hydrophobic | Amide-Pi Stacked |
|  |  | B:LYS48:NZ | 3.2 | UNL1:O | Hydrogen Bond | Conventional Hydrogen Bond |
|  |  | A:CYS61:N | 3 | UNL1:O | Hydrogen Bond | Conventional Hydrogen Bond |
|  |  | UNL1:H | 2.31 | A:CYS61:O | Hydrogen Bond | Conventional Hydrogen Bond |
|  |  | UNL1:H | 3.05 | B:SER50:OG | Hydrogen Bond | Conventional Hydrogen Bond |
|  |  | B:SER50:OG | 3 | UNL1:O | Hydrogen Bond | Conventional Hydrogen Bond |
|  |  | UNL1:H | 2.7 | A:LEU66:O | Hydrogen Bond | Conventional Hydrogen Bond |
|  |  | UNL1:C | 3.65 | A:GLU64:O | Hydrogen Bond | Conventional Hydrogen Bond |
|  |  | A:GLU64:CA | 3.37 | UNL1:F | Hydrogen Bond; Halogen | Carbon Hydrogen Bond;Halogen (Fluorine) |
|  |  | A:GLU64:OE2 | 3.34 | UNL1:F | Halogen | Halogen (Fluorine) |

**Figure 4.**
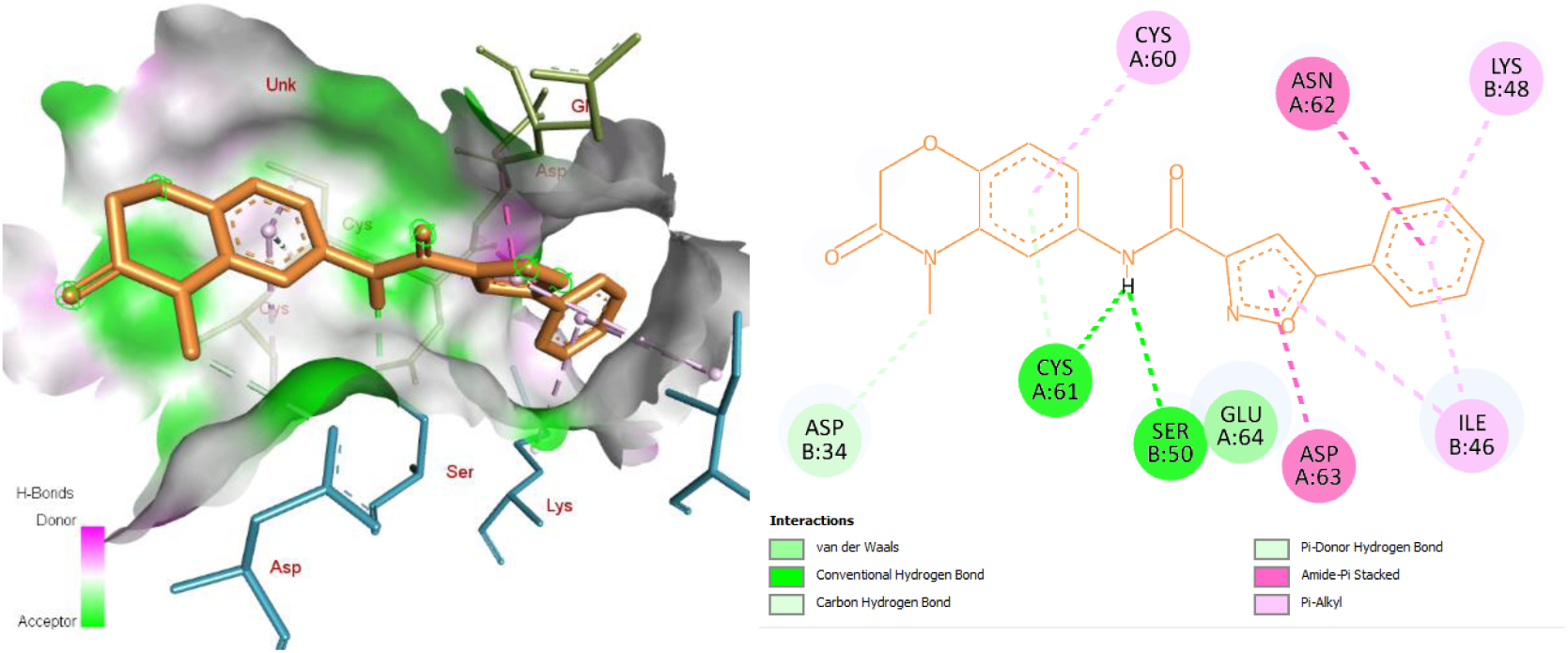
3D (left) and 2D (right) structure of the docked complex VEGFA and D519-0372. The violet and green color shades indicate hydrogen bonds.

**Figure 5.**
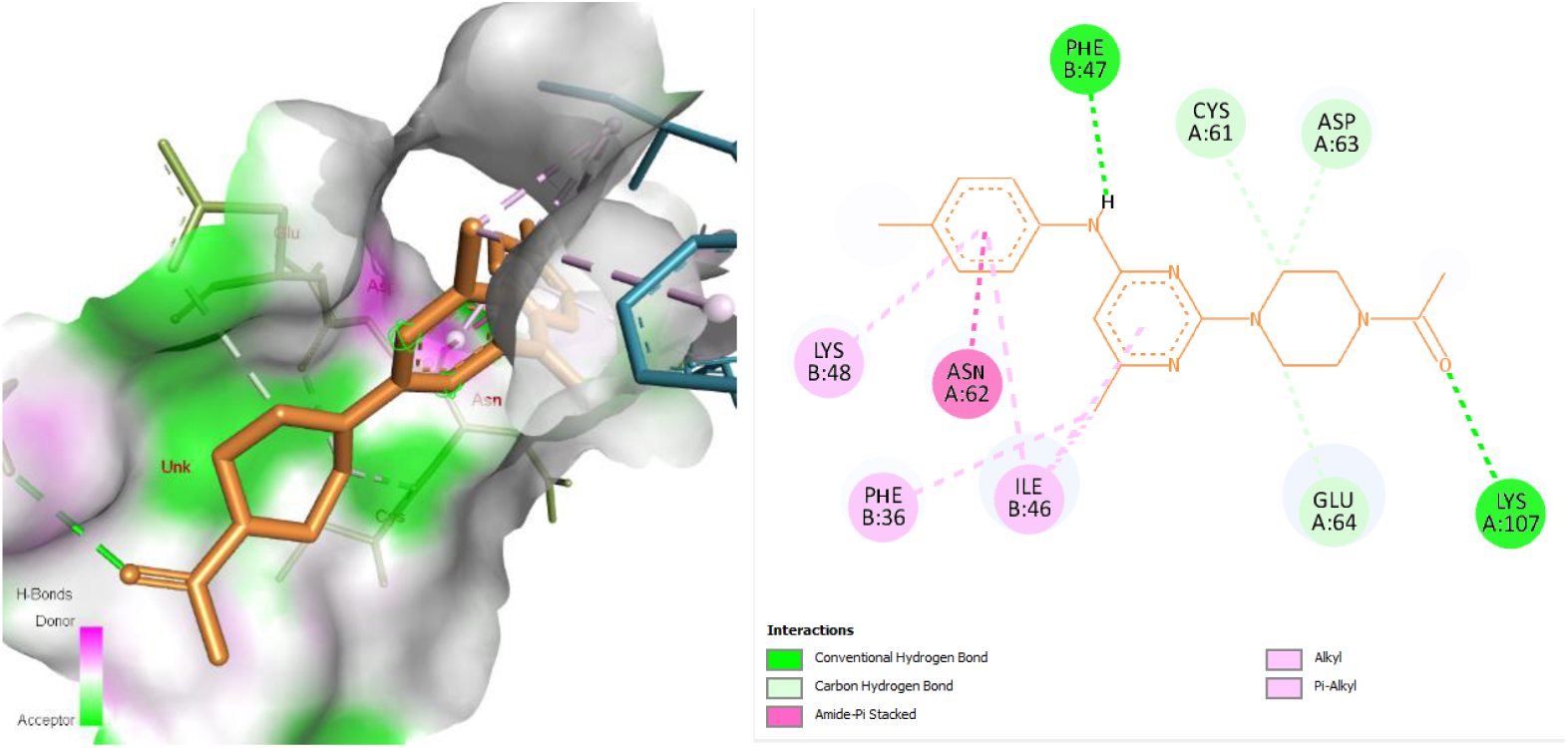
3D (left) and 2D (right) structure of the docked complex VEGFA and G868-0191. The violet and green color shades indicate hydrogen bonds.

**Figure 6.**
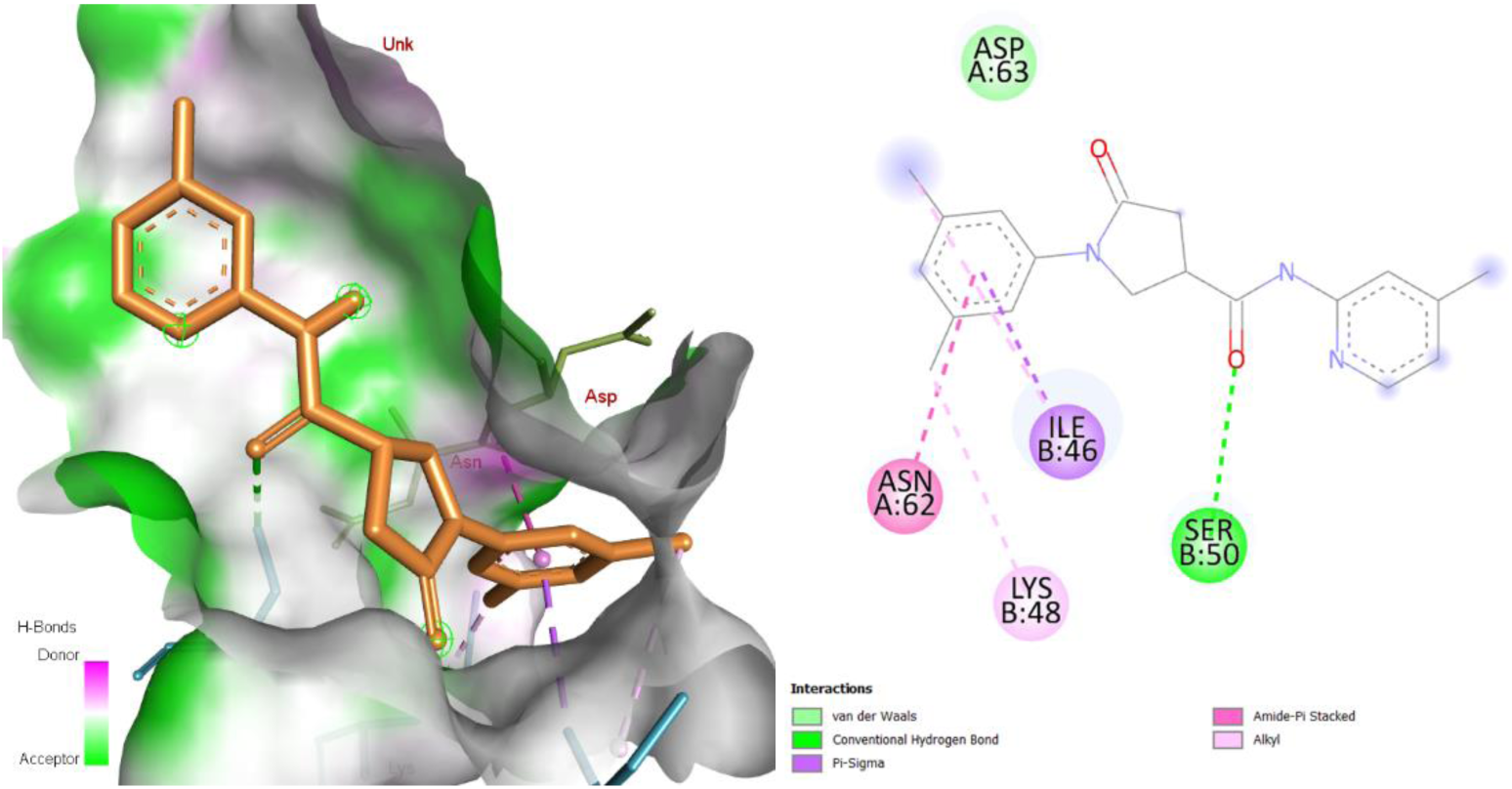
3D (left) and 2D (right) structure of the docked complex VEGFA and Y031-5201. The violet and green color shades indicate hydrogen bonds.

**Figure 7.**
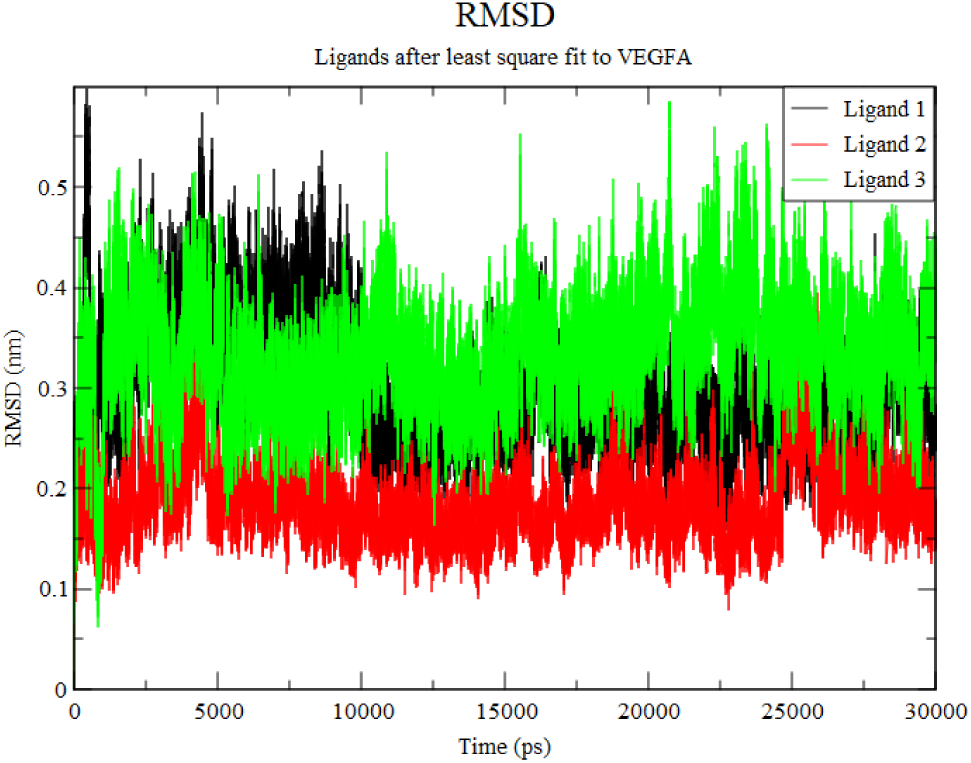
The plot displays the Root Mean Square Deviation (RMSD) values of three hit ligands from ChemDiv bound to the VEGFA protein, analyzed over a simulation time of 30 ns. Ligand order is the same as in Table I. Ligand 1 (Black): Exhibits moderate fluctuations throughout the simulation, with RMSD values generally ranging between 0.2 and 0.4 nm, indicating a relatively stable interaction with VEGFA. Ligand 2 (Red): Shows the lowest RMSD values overall, predominantly between 0.1 and 0.3 nm, suggesting the highest stability and potentially the strongest binding affinity to VEGFA among the three ligands. Ligand 3 (Green): Demonstrates higher fluctuations with RMSD values mostly between 0.3 and 0.5 nm, indicating less stability in its interaction with VEGFA compared to Ligand 2. This analysis suggests that Ligand 2 may have the most favorable binding characteristics with VEGFA, as indicated by its lower RMSD values, which reflect greater stability in the ligand-VEGFA complex over time.

**Figure 8.**
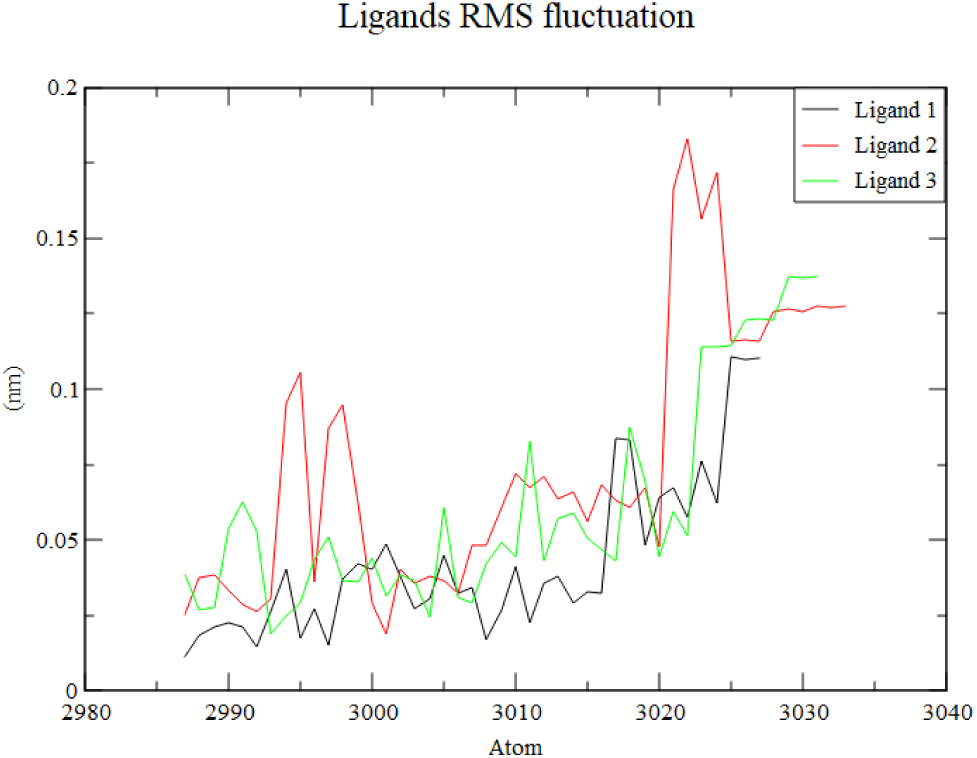
The plot shows the Root Mean Square Fluctuation (RMSF) of the three hit ligands from ChemDiv bound to VEGFA, as a function of the atom index along the ligand molecules. Ligand order is the same as in Table I. Ligand 1 (Black): Exhibits the lowest overall fluctuations, indicating that it maintains a relatively stable conformation when bound to VEGFA. Ligand 2 (Red): Displays higher fluctuations in certain regions, particularly around atom indices 3015-3025, suggesting areas of greater flexibility in the ligand structure, which might affect its binding stability. Ligand 3 (Green): Shows moderate fluctuations, with some peaks indicating areas of flexibility, though generally lower than those observed for Ligand 2. This analysis of RMSF provides insights into the conformational stability of the ligands when interacting with VEGFA, with Ligand 1 showing the most consistent and stable binding, while Ligand 2 exhibits regions of significant flexibility that could influence its interaction dynamics.

**Figure 9.**
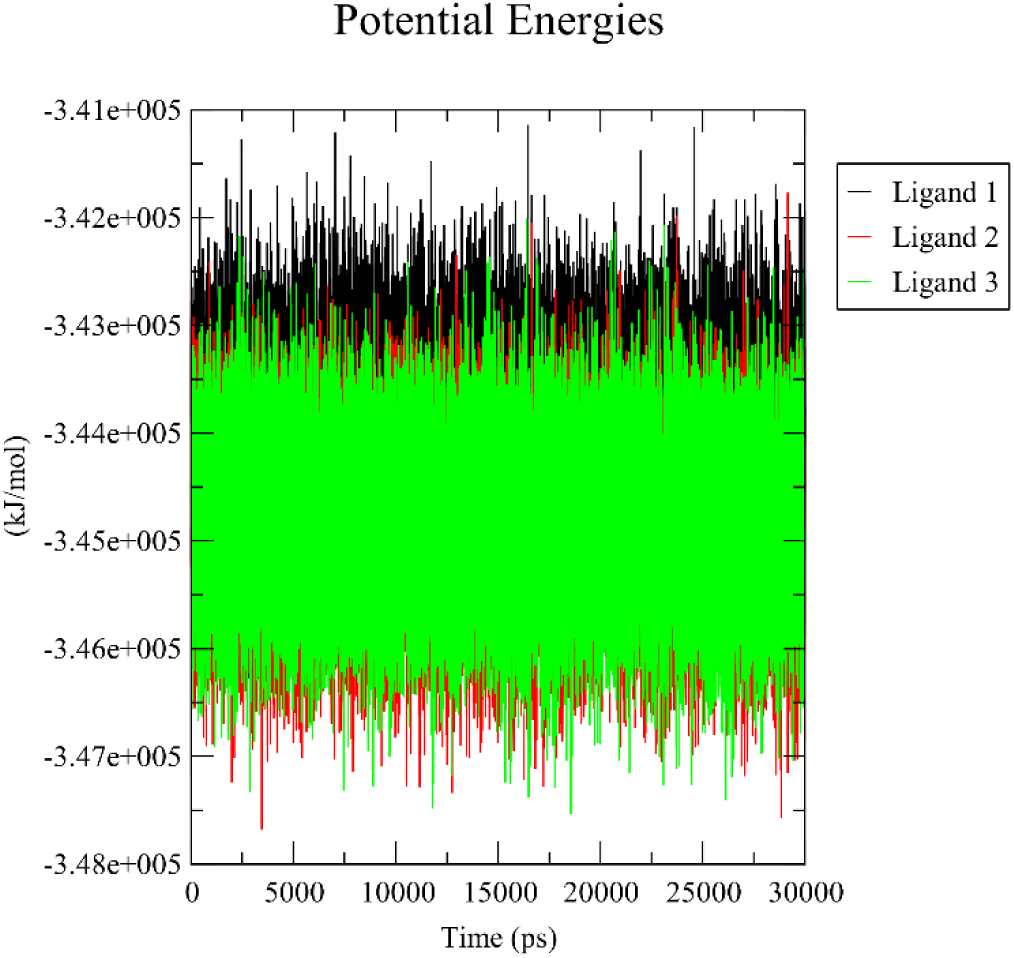
The plot depicts the potential energy profiles of the three hit ligands from ChemDiv bound to VEGFA during a molecular dynamics simulation spanning 30 ns. Ligand order is the same as in Table I. Ligand 1 (Black): Exhibits relatively higher potential energy values around −3.42 × 10^5 kJ/mol, suggesting that its interaction with VEGFA is less stable compared to the other ligands. Ligand 2 (Red): Shows slightly lower potential energy values, fluctuating around −3.45 × 10^5 kJ/mol, indicating a more stable binding interaction with VEGFA. Ligand 3 (Green): Demonstrates the lowest potential energy values, centered around −3.47 × 10^5 kJ/mol, suggesting the most stable interaction with VEGFA among the three ligands. Overall, Ligand 3 appears to form the most stable complex with VEGFA, as indicated by its consistently lower potential energy, while Ligand 1 shows the least stability.

**Figure 10.**
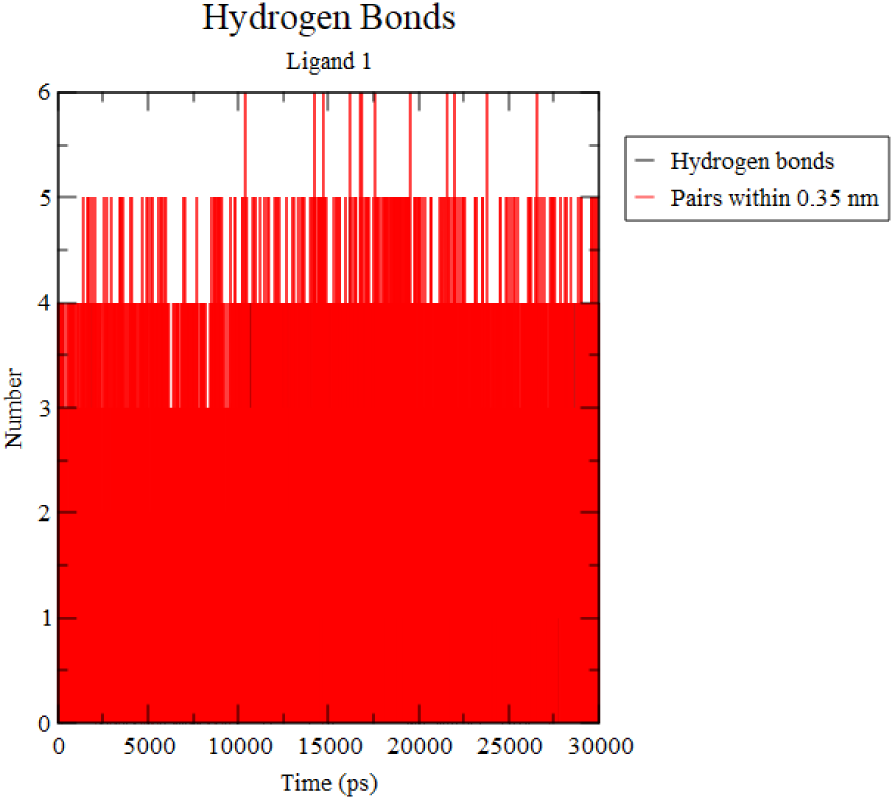
The plot illustrates the number of hydrogen bonds formed between Ligand 1 (D519-0372) and VEGFA during a 30 ns molecular dynamics simulation. The black line indicates the actual number of hydrogen bonds present between Ligand 1 and VEGFA at any given time during the simulation. The plot shows that the number of hydrogen bonds fluctuates between 2 and 5 throughout the simulation, with frequent formation of 3 to 4 hydrogen bonds. The red-shaded area represents the number of atom pairs within a distance of 0.35 nm, a typical cutoff distance for hydrogen bond formation. This indicates potential hydrogen bonds that might not meet all criteria (e.g., angle) for being classified as a hydrogen bond but are still in close proximity. The consistency in the number of hydrogen bonds, particularly around 3 to 4 bonds, suggests a stable interaction between Ligand 1 and VEGFA, which may contribute to the binding affinity and overall stability of the ligand-VEGFA complex.

**Figure 11.**
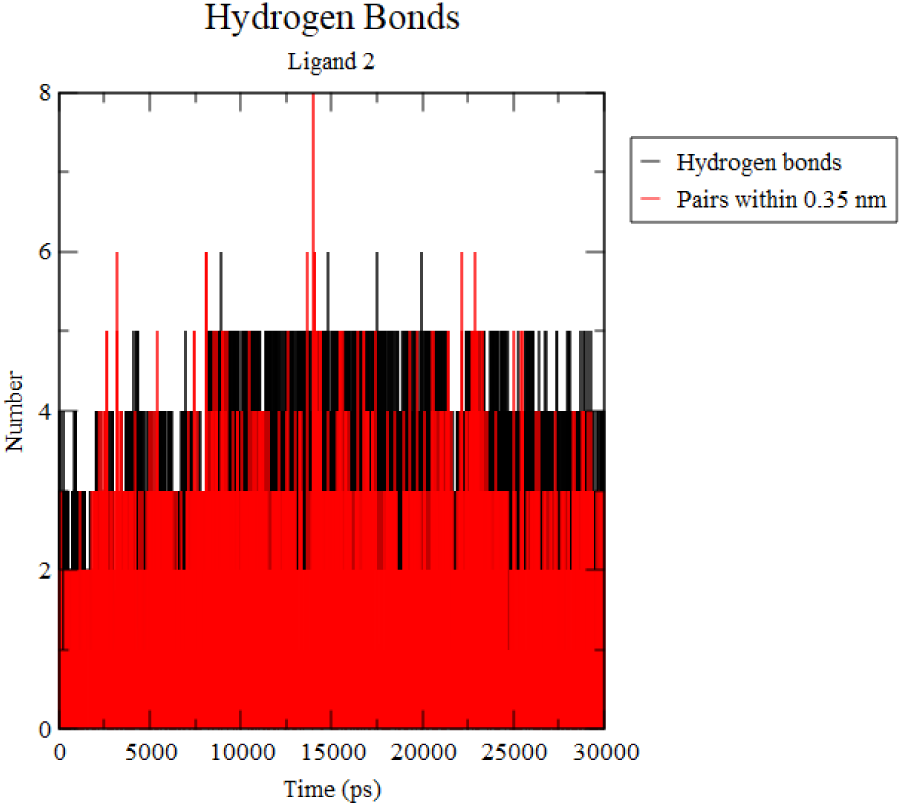
The plot shows the number of hydrogen bonds formed between Ligand 2 (G868-0191) and VEGFA during a 30 ns molecular dynamics simulation. The black line represents the number of hydrogen bonds between Ligand 2 and VEGFA at various time points during the simulation. The number of hydrogen bonds fluctuates between 2 and 6, with occasional peaks reaching up to 7 hydrogen bonds, indicating dynamic interactions between Ligand 2 and VEGFA. The red area represents the number of atom pairs within 0.35 nm, which is a typical distance cutoff for potential hydrogen bond formation. This region shows a consistent number of close contacts, suggesting possible hydrogen bonds that may not fully meet hydrogen bond criteria but are in proximity. The fluctuating number of hydrogen bonds, with frequent formation of 3 to 5 bonds, suggests that Ligand 2 has a more dynamic interaction with VEGFA, potentially forming and breaking hydrogen bonds more frequently. This dynamic behavior could influence the overall stability and binding affinity of Ligand 2 to VEGFA.

**Figure 12.**
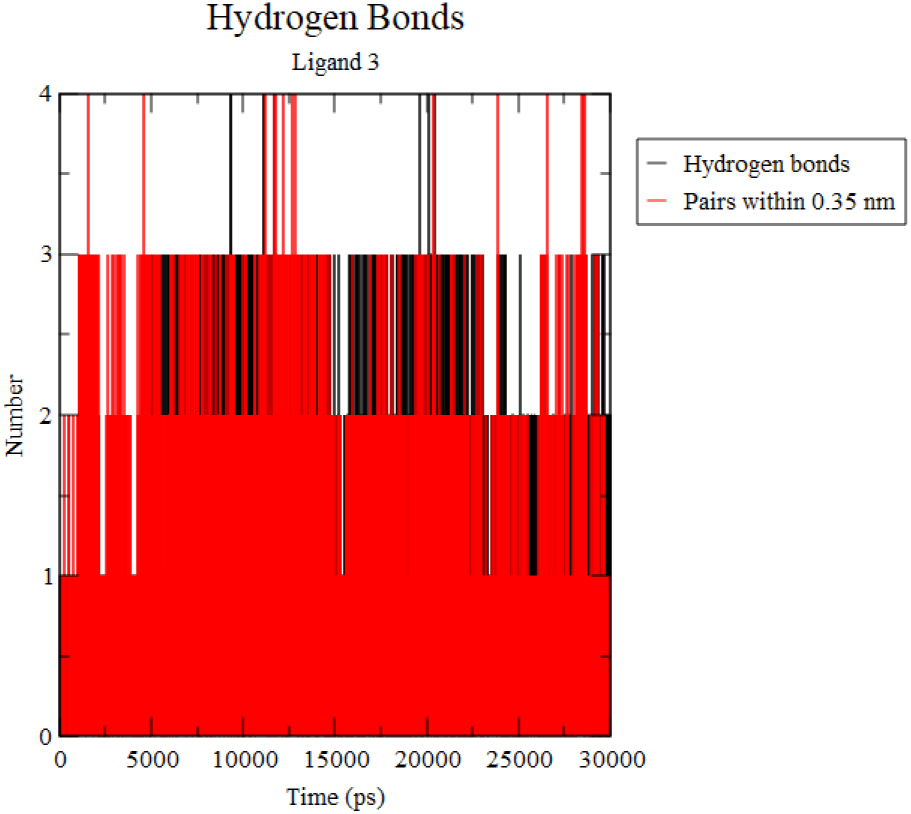
The plot illustrates the number of hydrogen bonds formed between Ligand 3 (Y031-5201) and VEGFA during a 30 ns molecular dynamics simulation. The black line shows the number of hydrogen bonds that Ligand 3 forms with VEGFA at different time points throughout the simulation. The number of hydrogen bonds fluctuates between 1 and 3, indicating that Ligand 3 generally maintains a lower number of hydrogen bonds with VEGFA compared to the other ligands. The red-shaded area represents the number of atom pairs within a 0.35 nm distance, which suggests potential hydrogen bonding interactions. The plot shows a consistent proximity of atoms, with occasional formation of hydrogen bonds. The relatively low number of hydrogen bonds, often ranging between 1 and 2, suggests that Ligand 3 forms weaker or less frequent hydrogen bonds with VEGFA. This could imply a less stable interaction compared to Ligands 1 and 2, potentially affecting the overall binding affinity of Ligand 3 to VEGFA.

**Figure 13.**
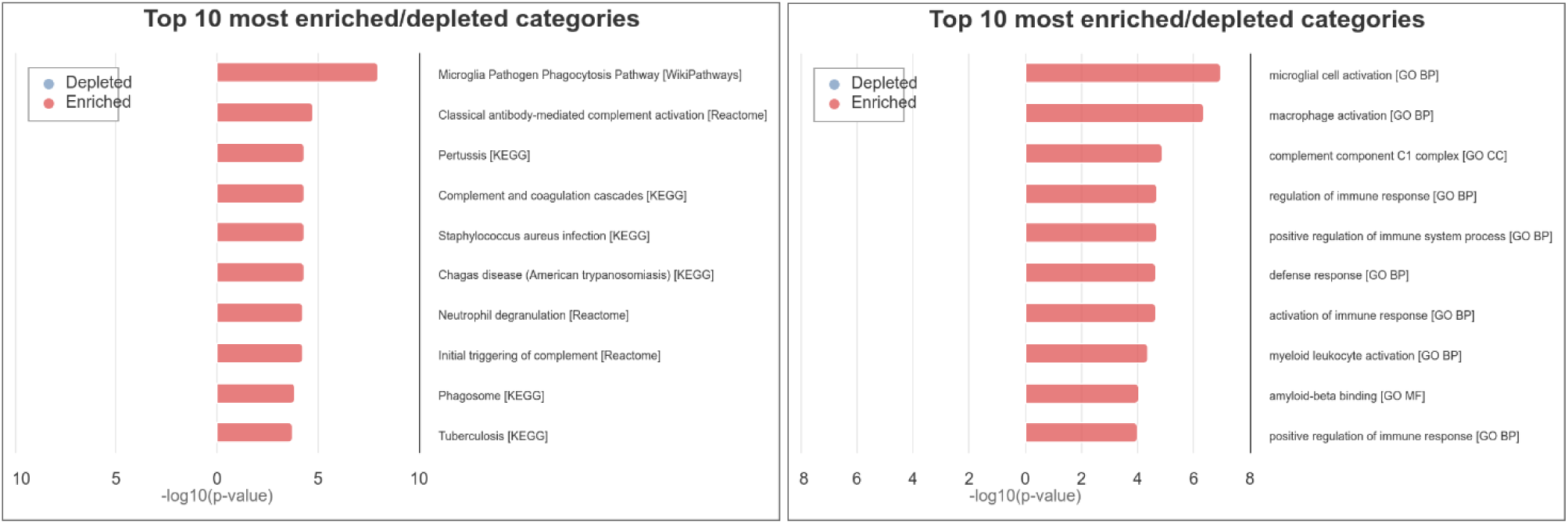
Top 10 enriched categories for pathways (left) and gene ontology (right) for upregulated genes in glioblastomas. The left plot illustrates the most enriched biological pathways related to up-regulated genes in glioblastomas. These include immune-related processes such as the “Microglia Pathogen Phagocytosis Pathway” and “Classical antibody-mediated complement activation,” as well as pathways related to infections, complement activation, and phagosome functions (KEGG and Reactome databases). The right plot highlights the most enriched Gene Ontology (GO) categories, focusing on immune activation and response. The most enriched GO categories include “Microglial cell activation,” “Macrophage activation,” “Complement component C1 complex,” and various immune system processes. This suggests that up-regulated genes are involved in modulating immune responses within the glioblastoma microenvironment.

**Figure 14.**
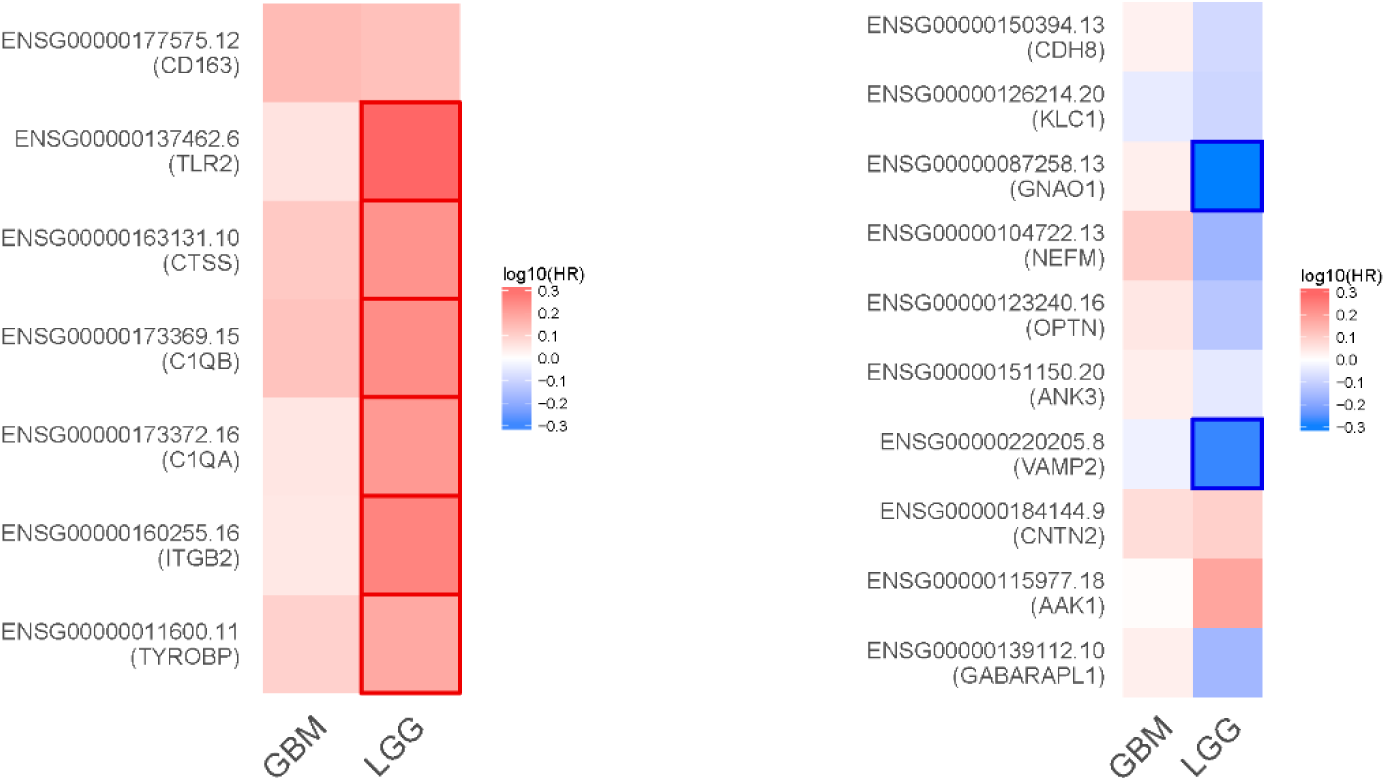
Survival analysis of upregulated and down-regulated hub genes in glioblastomas (GBM) and low-grade gliomas (LGG). The left heatmap shows the log10-transformed hazard ratios (HR) for up-regulated hub genes across GBM and LGG. Genes like CD163, TLR2, and C1QA exhibit higher expression in LGG, correlating with worse survival outcomes, as indicated by positive log10(HR) values (red). The right heatmap represents the survival analysis for down-regulated hub genes across the same tumor types. Genes such as KLC1, GNAO1, and VAMP2 display lower expression in LGG, correlating with better survival outcomes, as indicated by negative log10(HR) values (blue). These analyses highlight potential gene expression patterns that influence survival in glioblastomas and low-grade gliomas, providing insight into tumor biology and potential therapeutic targets.

## Discussion

### Novel VEGFA inhibitors

Hits from the ChemDiv library exhibited superior scores and represent novel compounds not documented in existing literature, indicating their potential for lead development. The stability and effectiveness of these inhibitors were further supported by the molecular dynamics simulation results, where Ligand 2 (G868-0191), in particular, demonstrated the highest stability in its interaction with VEGFA, as indicated by the RMSD, RMSF, and potential energy analyses, further increasing their potential for drug development. Despite VEGFA not being a kinase, the screening of kinase inhibitors was based on an in-silico study demonstrating binding affinity between VEGFA and tyrosine kinase inhibitors [67], a hypothesis supported by the current findings.

The hits from the ZINC database also show potential for drug development. They have good toxicity and druglikeness properties, just behind the hits from ChemDiv, but also outperforming control drugs. They consist of flavonoids, with 5,4’-Dihydroxyflavone and Liquiritigenin classified as flavones, and 4’-Hydroxyflavanone identified as a flavanone. The analysis of the docked complexes revealed a substantial number of Pi interactions, suggesting improved structural stability and specificity, since these interactions help to anchor the ligand in the binding pocket, enhancing its stability. Also, the arrangement and orientation of aromatic rings in both the ligand and the protein can create a unique interaction pattern that contributes to selective binding [68–71]. Previous studies have reported these compounds as inhibitors of tankyrase [72] and monoamine oxidase [73] (4’-Hydroxyflavanone), androgen receptor [74] (5,4’-Dihydroxyflavone), and aromatase [75] (Liquiritigenin), which hint to a good bioavailability, although their potential as VEGFA inhibitors remains unexplored.

Our study identified two potential candidates for drug repurposing: sunitinib and ticlopidine hydrochloride. Sunitinib, a multi-targeted receptor tyrosine kinase inhibitor, demonstrated the highest score in our analysis, surpassing the initial hits. However, its potential is somewhat diminished by a Brenk alert and its identification as an irritant in toxicity screenings. These factors could complicate its use in clinical applications, potentially requiring careful administration or novel delivery methods to mitigate these issues. Despite these challenges, Sunitinib’s ability to cross the blood-brain barrier and its broad-spectrum activity against multiple oncogenic pathways still make it a candidate worth considering for further investigation.

In contrast, ticlopidine hydrochloride emerged as a more promising candidate for repurposing. This antiplatelet drug, traditionally used to reduce the risk of stroke and heart attack, exhibited no significant drawbacks in our analysis. Its ranking between the hits from ChemDiv and ZINC databases, coupled with its well-established safety profile, positions it as a particularly attractive option for repurposing. The lack of identified limitations in our study suggests that ticlopidine hydrochloride could potentially transition more smoothly into clinical trials for new indications. However, further research is needed to fully elucidate its mechanism of action in the context of cancer treatment and to determine optimal dosing strategies that may differ from its current therapeutic use

While Irinotecan was the highest-ranked control drug, it exhibited Lipinski and Ghose violations, specifically it has a molecular weight greater than 480 g/mol (Lipinski and Ghose), a molar refractivity greater than 130, and more than 70 atoms (Ghose). Besides, none of the controls passed the blood-brain barrier test (boiled egg method in SwissADME), hindering their use as glioblastomas drugs.

The identified VEGFA inhibitors, including repurposed drugs, demonstrate promising potential as therapeutic agents. These inhibitors could disrupt the VEGFA signaling pathway, thereby impeding the vascularization of tumors, which is critical for their growth and survival. Their integration into current glioblastoma treatment protocols presents a promising avenue for improving patient outcomes. These inhibitors could be incorporated at various stages of the treatment process, including in combination with the standard Stupp protocol (maximal safe resection, radiotherapy, and temozolomide), as neoadjuvant therapy to facilitate surgical resection, or as maintenance therapy to delay recurrence [76]. For recurrent glioblastoma, these inhibitors might serve as monotherapy or in combination with other salvage treatments. Moreover, their potential synergy with emerging immunotherapies warrants investigation, given VEGF’s known immunosuppressive effects in the tumor microenvironment [77].

The implementation of this novel VEGFA inhibitors would likely follow a personalized approach, based on molecular profiling of individual tumors to identify patients most likely to benefit [78]. Integration strategies may also explore local delivery methods to maximize efficacy while minimizing systemic side effects. Importantly, these new inhibitors could offer alternatives for patients who develop resistance to current anti-VEGF therapies like bevacizumab [79]. However, successful integration will depend on rigorous clinical trials to establish efficacy, safety, and optimal combination strategies with existing treatments.

Ultimately, the goal of integrating novel VEGFA inhibitors into glioblastoma treatment protocols is to improve survival outcomes and quality of life for patients. This may be achieved through more durable responses, delayed recurrence, or better control of symptoms related to tumor growth and edema. As research progresses, it will be crucial to carefully consider the timing, dosing, and combination of these inhibitors with other treatments to maximize benefits while managing potential side effects and drug interactions [80].

### Hub genes as therapeutic targets and biomarkers

The analysis of hub genes in glioblastomas reveals two distinct patterns: the overexpression of immune-related genes and the downregulation of genes associated with neuronal and synaptic functions. Hub genes TYROBP, ITGB2, C1QA, C1QB, CTSS, TLR2, and CD163 are significantly overexpressed in glioblastomas. These genes are primarily involved in immune response pathways, including macrophage activation, complement system regulation, and inflammatory signaling. Their overexpression suggests an immunosuppressive and pro-inflammatory tumor microenvironment that facilitates glioblastoma growth and progression [81,83]. Specifically, markers like CD163 and TLR2 are associated with the M2 macrophage phenotype, promoting immune evasion, and contributing to poorer survival outcomes [82]. Targeting these immune-related pathways could present novel therapeutic opportunities, such as reprogramming tumor-associated macrophages to counteract the tumor-supportive microenvironment [83].

Conversely, hub genes GABARAPL1, AAK1, CNTN2, VAMP2, ANK3, OPTN, NEFM, GNAO1, KLC1, and CDH8 are significantly downregulated in glioblastomas. These genes participate in key neuronal processes such as autophagy, synaptic function, cytoskeletal organization, and cellular adhesion. Their downregulation reflects a loss of normal neuronal characteristics and cellular communication, which may contribute to the aggressive, invasive nature of glioblastomas [84,85]. This pattern suggests that the down-regulated genes are not only involved in the loss of tumor suppressive functions but could also serve as valuable biomarkers for glioblastoma diagnosis or prognosis [85]. For example, genes such as GABARAPL1 and OPTN, which participate in autophagy [86], and CDH8, involved in cell adhesion [87], could be explored as potential biomarkers for early detection or as indicators of tumor progression. The disruption of autophagy and cell adhesion mechanisms correlates with tumor aggressiveness, and their expression levels may provide insights into tumor behavior and response to therapy.

In conclusion, these findings emphasize the dual nature of glioblastoma biology, driven by an overactive immune component and a loss of neuronal functionality. Targeting the immune microenvironment, while leveraging down-regulated genes as biomarkers, holds promise for developing more personalized therapeutic strategies and improving patient outcomes in glioblastomas.

### Enrichment Analysis and Importance of Identified Hub Genes TYROBP and ITGB2

The enrichment analysis highlights several key pathways and biological processes associated with up-regulated genes in glioblastomas, particularly those involved in immune responses. Pathways such as “Microglia Pathogen Phagocytosis Pathway,” “Classical antibody-mediated complement activation,” and “Complement and coagulation cascades” underscore the critical role of immune and inflammatory processes in glioblastoma progression [88,89]. The overrepresentation of immune-related pathways aligns with the findings from gene expression analyses, where genes like TYROBP and ITGB2 were identified as hub genes that are significantly up-regulated in glioblastomas.

TYROBP (also known as DAP12) is a key regulator of immune signaling, particularly in microglia, macrophages, and other myeloid cells [90]. It plays an essential role in activating immune responses and mediating phagocytosis, both of which are crucial in the glioblastoma microenvironment. The enrichment of microglial activation pathways suggests that glioblastomas exploit the tumor-supportive properties of microglia, leading to a pro-tumorigenic, immunosuppressive environment that promotes tumor growth and immune evasion [88,91]. Targeting TYROBP may offer therapeutic potential by disrupting these tumor-promoting signals, reprogramming the immune response to inhibit tumor progression [92].

ITGB2 (CD18), a critical integrin involved in immune cell adhesion and signaling, is another important hub gene identified in glioblastomas. Its overexpression is linked to enhanced recruitment and activation of immune cells, particularly tumor-associated macrophages (TAMs), which are known to adopt an M2-like, tumor-supportive phenotype in glioblastomas [93]. ITGB2 interacts with various complement and coagulation pathways, further supporting the relevance of the enrichment analysis. The significant enrichment of complement activation pathways, combined with the up regulation of ITGB2, points to a key role for the complement system in modulating immune responses within the tumor microenvironment [94]. Therapeutically targeting ITGB2 could impair the tumor-promoting functions of TAMs and other immune cells, thereby shifting the balance towards anti-tumor immunity [95].

In summary, the enrichment analysis not only reinforces the biological significance of immune-related processes in glioblastoma progression but also underscores the critical roles of TYROBP and ITGB2 as hub genes. These findings suggest that these genes could serve as potential therapeutic targets to modulate the immune microenvironment in glioblastomas, offering new avenues for the development of immunotherapy strategies [93,95].

### Combining Databases to Address Tumor Heterogeneity

Glioblastomas are characterized by high inter- and intra-tumor heterogeneity, which can vary significantly across different populations and subtypes. A single dataset or source may not fully capture the complexity of gene expression patterns in diverse glioblastoma cases. By combining gene expression data from both the GEO and GEPIA2 databases, our approach integrates multiple cohorts and tumor profiles, helping to overcome limitations related to population bias and sample variability. The use of GEO datasets allows us to capture expression patterns from two independent studies, each potentially representing different subpopulations, while GEPIA2 integrates data from The Cancer Genome Atlas (TCGA) and GTEx databases. This combined approach increases the robustness of our findings by identifying common up-regulated and down-regulated hub genes across multiple datasets, thus reflecting broader population diversity. This strategy ensures that the hub genes identified are likely relevant across a wider spectrum of glioblastoma cases, enhancing their potential for general clinical application. Furthermore, the use of two metrics, interaction degree and betweenness, to obtain the hub genes, increases the reliability and robustness of the findings.

### Study Limitations

While our study provides valuable insights into potential therapeutic targets and biomarkers in glioblastomas, it is important to acknowledge the limitations of purely in silico approaches. Computational analyses, such as those performed here, rely on existing databases and algorithms to predict gene interactions, regulatory pathways, and potential drug targets. These predictions are inherently limited by the quality and completeness of the available data and can sometimes overlook important biological nuances, such as post-translational modifications or tumor microenvironment interactions, which may not be fully captured in gene expression data. Moreover, the absence of experimental validation means that the observed patterns of overexpression or down-regulation may not always directly translate into functional outcomes in vivo. Therefore, while bioinformatics is a powerful tool for hypothesis generation, its findings must be interpreted cautiously and validated through experimental studies before clinical application.

### Perspectives and Future Directions

Moving forward, the hub genes identified in this study offer promising avenues for further research, particularly as potential therapeutic targets, and biomarkers in glioblastomas. However, to translate these findings into clinical practice, extensive in vitro and in vivo validation is necessary. In vitro experiments, such as CRISPR/Cas9 gene editing or RNA interference, could be employed to knock down or overexpress the identified hub genes in glioblastoma cell lines to confirm their role in tumor proliferation, invasion, and immune modulation. Additionally, 3D organoid models and co-culture systems could be used to mimic the tumor microenvironment and assess the functional consequences of targeting immune-related hub genes, such as CD163 and TLR2, which are associated with immunosuppressive phenotypes. In vivo studies using glioblastoma mouse models will be crucial for validating the therapeutic potential of these hub genes and assessing the efficacy of targeting them with small molecule inhibitors or antibodies.

Furthermore, the down-regulated hub genes, particularly GABARAPL1, OPTN, and CDH8, could be explored as biomarkers for glioblastoma diagnosis or prognosis in clinical settings. Liquid biopsy approaches, such as circulating tumor DNA (ctDNA) or RNA (ctRNA) analyses, could help determine whether these genes are detectable in the blood or cerebrospinal fluid, providing a non-invasive means of early detection or monitoring disease progression. Ultimately, integrating these findings into preclinical and clinical research pipelines will be essential to fully realize the therapeutic and diagnostic potential of these hub genes.

In conclusion, the combined in silico approach has provided significant insights into glioblastoma biology, but further validation is critical to harness the clinical utility of the identified targets and biomarkers.

## Conclusions

This study provides a comprehensive analysis of gene expression data from GEO studies and GEPIA2, identifying key hub genes such as TYROBP, ITGB2, C1QA, C1QB, CTSS, TLR2, and CD163 that are significantly overexpressed in glioblastomas, with a prominent role in immune regulation. Our enrichment analysis further reveals that these genes participate in crucial immune and inflammatory pathways, emphasizing the immune-suppressive and tumor-promoting microenvironment of glioblastomas. The downregulation of key neuronal and synaptic function genes such as GABARAPL1, OPTN, and CDH8 suggests their potential use as biomarkers for early detection or prognosis.

The computational discovery of VEGFA inhibitors offers promising leads for drug development, with hits from the ChemDiv and ZINC databases outperforming current control drugs in binding stability and druglikeness properties. In particular, G868-0191 showed the highest stability in molecular dynamics simulations, making it a potential candidate for further experimental validation. The identification of two repurposed drugs, Sunitinib and Ticlopidine hydrochloride, adds to the therapeutic landscape, though further research is required to confirm their efficacy and safety in clinical settings.

In conclusion, the integration of novel VEGFA inhibitors into glioblastoma treatment protocols, alongside the identification of hub genes as potential therapeutic targets and biomarkers, provides a multifaceted approach to improving patient outcomes. Future research must focus on in vitro and in vivo validation to ensure the clinical applicability of these findings.

**Authors declare no conflict of interest. Ethical approval was not required since no animals or humans were used as subjects in the present study.**

## Authors’ contribution

**Angélica Bautista**

Conceptualization (Supporting), Data curation (Lead), Formal analysis (Equal), Investigation (Equal), Methodology (Equal), Project administration (Supporting), Validation (Equal), Writing – original draft (Equal), Writing – review & editing (Equal).

**Ricardo Romero**

Conceptualization (Lead), Data curation (Supporting), Formal analysis (Equal), Investigation (Equal), Methodology (Equal), Project administration (Lead), Supervision (Lead), Validation (Equal), Writing – original draft (Equal), Writing – review & editing (Equal).

## Acknowledgements

Angélica Bautista thanks Universidad Autónoma Metropolitana for granting her a scholarship to conduct her social service, from which the current work arose. Authors declare that the present study was not funded by any public or private agency.

## References

1. Zong H, Parada LF, Baker SJ. Cell of origin for malignant gliomas and its implication in therapeutic development. Cold Spring Harb Perspect Biol. 2015;7(5):1–13.

2. Louis DN, Perry A, Reifenberger G, et al. The 2016 World Health Organization Classification of Tumors of the Central Nervous System: a summary. Acta Neuropathol. 2016;131(6):803–820. doi:10.1007/s00401-016-1545-1

3. Reifenberger G, Wirsching HG, Knobbe-Thomsen CB, Weller M. Advances in the molecular genetics of gliomas-implications for classification and therapy. Nat Rev Clin Oncol. 2017;14(7):434–52.

4. Gimple RC, Bhargava S, Dixit D, Rich JN. Glioblastoma stem cells: Lessons from the tumor hierarchy in a lethal cancer. Genes Dev. 2019;33(11–12):591–609.

5. Ostrom QT, Patil N, Cioffi G, et al. CBTRUS Statistical Report: Primary Brain and Other Central Nervous System Tumors Diagnosed in the United States in 2013-2017. Neuro Oncol. 2020;22(12 Suppl 2):iv1–iv96. doi:10.1093/neuonc/noaa200.

6. Ghiaseddin AP, Shin D, Melnick K, Tran DD. Tumor Treating Fields in the Management of Patients with Malignant Gliomas. Curr Treat Options Oncol. 2020;21(9).

7. Wirsching HG, Galanis E, Weller M. Glioblastoma. Handb Clin Neurol. 2016; 134:381–97.

8. Omuro A, DeAngelis LM. Glioblastoma and other malignant gliomas: A clinical review. JAMA. 2013;310(17):1842–50.

9. Sun T, Warrington NM, Luo J, et al. Sexually dimorphic RB inactivation underlies mesenchymal glioblastoma prevalence in males. J Clin Invest. 2014;124(9):4123–4133. doi:10.1172/JCI71048

10. Bondy ML, Scheurer ME, Malmer B, et al. Brain tumor epidemiology: consensus from the Brain Tumor Epidemiology Consortium. Cancer. 2008;113(7 Suppl):1953–1968. doi:10.1002/cncr.23741

11. Butowski NA. Epidemiology and diagnosis of brain tumors. Continuum (Minneap Minn). 2015;21(2 Neuro-oncology):301–313. doi:10.1212/01.CON.0000464171.50638.fa

12. Plate KH, Breier G, Weich HA, Risau W. Vascular endothelial growth factor is a potential tumour angiogenesis factor in human gliomas in vivo. Nature. 1992;359(6398):845–848. doi:10.1038/359845a0

13. Alves B, Peixoto J, Macedo S, Pinheiro J, Carvalho B, Soares P, et al. High VEGFA Expression Is Associated with Improved Progression-Free Survival after Bevacizumab Treatment in Recurrent Glioblastoma. Cancers (Basel). 2023;15(8).

14. Holst CB, et al. Perspective: targeting VEGF-A and YKL-40 in glioblastoma–matter matters. Cell Cycle [Internet]. 2021;20(7):702–15. 10.1080/15384101.2021.1901037

15. Goel, H. L., & Mercurio, A. M. (2013). VEGF targets the tumour cell. Nature Reviews Cancer, 13(12), 871–882.

16. Olsson, A. K., Dimberg, A., Kreuger, J., & Claesson-Welsh, L. (2006). VEGF receptor signalling—in control of vascular function. Nature Reviews Molecular Cell Biology, 7(5), 359–371.

17. Apte, R. S., Chen, D. S., & Ferrara, N. (2019). VEGF in signaling and disease: beyond discovery and development. Cell, 176(6), 1248–1264.

18. Tugues, S., Koch, S., Gualandi, L., Li, X., & Claesson-Welsh, L. (2011). Vascular endothelial growth factors and receptors: anti-angiogenic therapy in the treatment of cancer. Molecular Aspects of Medicine, 32(2), 88–111.

19. Ferrara, N., & Adamis, A. P. (2016). Ten years of anti-vascular endothelial growth factor therapy. Nature Reviews Drug Discovery, 15(6), 385–403.

20. Friedman HS, Prados MD, Wen PY, et al. Bevacizumab alone and in combination with irinotecan in recurrent glioblastoma. J Clin Oncol. 2009;27(28):4733–4740. doi:10.1200/JCO.2008.19.8721

21. Lu KV, Chang JP, Parachoniak CA, et al. VEGF inhibits tumor cell invasion and mesenchymal transition through a MET/VEGFR2 complex. Cancer Cell. 2012;22(1):21–35. doi:10.1016/j.ccr.2012.05.037

22. Cloughesy TF, Cavenee WK, Mischel PS. Glioblastoma: from molecular pathology to targeted treatment. *Annu Rev Pathol*. 2014;9:1–25. doi:10.1146/annurev-pathol-011110-130324

23. Cloughesy TF, Chang SM, Wen PY, et al. Phase III trial of bevacizumab added to standard-of-care treatment for glioblastoma: final results of the AVAglio trial. *J Clin Oncol*. 2014;32(22):2342–2349. doi:10.1200/JCO.2013.53.3870

24. Stanzione F, Giangreco I, Cole JC. Use of molecular docking computational tools in drug discovery [Internet]. 1st ed. Vol. 60, Progress in Medicinal Chemistry. Elsevier B.V.; 2021. 273–343 p. 10.1016/bs.pmch.2021.01.004

25. Shaker B, Ahmad S, Lee J, Jung C, Na D. In silico methods and tools for drug discovery. Comput Biol Med [Internet]. 2021;137(September):104851. 10.1016/j.compbiomed.2021.104851

26. Li K, Du Y, Li L, Wei DQ. Bioinformatics Approaches for Anti-cancer Drug Discovery. Curr Drug Targets. 2019;21(1):3–17.

27. Chen H, Engkvist O, Wang Y, Olivecrona M, Blaschke T. The rise of deep learning in drug discovery. Drug Discov Today. 2018;23(6):1241–1250. doi:10.1016/j.drudis.2018.01.039

28. Cancer Genome Atlas Research Network. Comprehensive genomic characterization defines human glioblastoma genes and core pathways. Nature. 2008;455(7216):1061–1068. doi:10.1038/nature07385

29. Grzmil M, Morin P Jr, Lino MM, Merlo A et al. MAP kinase-interacting kinase 1 regulates SMAD2-dependent TGF-β signaling pathway in human glioblastoma. Cancer Res 2011 Mar 15;71(6):2392–402. PMID: 21406405

30. Sun L, Hui AM, Su Q, Vortmeyer A et al. Neuronal and glioma-derived stem cell factor induces angiogenesis within the brain. Cancer Cell 2006 Apr;9(4):287–300. PMID: 16616334

31. Fonseka, P., Pathan, M., Chitti, S.V., Kang, T. and Mathivanan, S. (2021) FunRich enables enrichment analysis of OMICs datasets. Journal of Molecular Biology. 166747.

32. Pathan, M., Keerthikumar, et al. A novel community driven software for functional enrichment analysis of extracellular vesicles data. J Extracellular Vesicles. 2017. 1:1321455.

33. Pathan, M., et al. FunRich: a standalone tool for functional enrichment analysis. Proteomics. 2015. 15, 2597–2601.

34. Tang, Z. et al. (2019) GEPIA2: an enhanced web server for large-scale expression profiling and interactive analysis. Nucleic Acids Res, 10.1093/nar/gkz430.

35. The GTEx Consortium, The GTEx Consortium atlas of genetic regulatory effects across human tissues. Science 369,1318–1330(2020). DOI:10.1126/science.aaz1776

36. Szklarczyk D., et al. The STRING database in 2023: protein–protein association networks and functional enrichment analyses for any sequenced genome of interest. Nucleic Acids Res. 2023 Jan 6;51(D1): D638–646.

37. Szklarczyk D., et al. The STRING database in 2021: customizable protein–protein networks, and functional characterization of user-uploaded gene/measurement sets. Nucleic Acids Res. 2021 Jan 8;49(D1): D605–12.

38. Chin, Chia-Hao et al. “cytoHubba: identifying hub objects and sub-networks from complex interactome.” BMC systems biology vol. 8 Suppl 4, Suppl 4. 2014: S11. 10.1186/1752-0509-8-S4-S11.

39. Shannon, Paul, et al. “Cytoscape: a software environment for integrated models of biomolecular interaction networks.” Genome research vol. 13,11. 2003: 2498–504. 10.1101/gr.1239303

40. Muller YA, et al. Vascular endothelial growth factor: crystal structure and functional mapping of the kinase domain receptor binding site. Proc Natl Acad Sci U S A. 1997;94(14):7192–7197. 10.1073/pnas.94.14.7192

41. Pettersen EF, et al. UCSF Chimera, A visualization system for exploratory research and analysis. Journal of Computational Chemistry. 2004;25(13):1605–12. 10.1002/jcc.20084

42. Tian, et al. Nucleic Acids Res. 2018. PMID: 29860391 DOI: 10.1093/nar/gky473.

43. Guex N, Peitsch MC. SWISS-MODEL and the Swiss-Pdb Viewer: An environment for comparative protein modeling. ELECTROPHORESIS. 1997;18(15):2714–23. 10.1002/elps.1150181505

44. Christen M, Hünenberger PH, Bakowies D, et al (2005) The GROMOS software for biomolecular simulation: GROMOS05. J. Comput. Chem. 26:1719–1751

45. ChemDiv. Protein Kinases Inhibitors Library; 2024. https://www.chemdiv.com/catalog/focused-and-targeted-libraries/protein-kinases-inhibitors-library

46. O’Boyle NM, et al. Open Babel: An open chemical toolbox. Journal of Cheminformatics. 2011 Oct; 3(1):33. 10.1186/1758-2946-3-33.

47. Trott O, Olson AJ. AutoDock Vina: Improving the speed and accuracy of docking with a new scoring function, efficient optimization, and multithreading. Journal of Computational Chemistry. 2010;31(2):455–61. 10.1002/jcc.21334.

48. Eberhardt J, Santos-Martins D, Tillack AF, Forli S. AutoDock Vina 1.2.0: New Docking Methods, Expanded Force Field, and Python Bindings. Journal of Chemical Information and Modeling. 2021 Aug;61(8):3891–8. 10.1021/acs.jcim.1c00203.

49. Daina A, Michielin O, Zoete V. Swiss ADME: a free web tool to evaluate pharmacokinetics, drug-likeness and medicinal chemistry friend-liness of small molecules. Scientific Reports. 2017 Mar;7(1):42717. 10.1038/srep42717.

50. Sander T, Freyss J, von Korff M, Reich JR, Rufener C. OSIRIS, an entirely in-house developed drug discovery informatics system. J Chem Inf Model. 2009;49(2):232–246. 10.1021/ci800305f.

51. Lipinski CA, Lombardo F, Dominy BW, Feeney PJ. Experimental and computational approaches to estimate solubility and permeability in drug discovery and development. The article was originally published in Advanced Drug Delivery Reviews 23 (1997) 3–25.1. 10.1016/S0169-409X(00)00129-0.

52. Ghose AK, Viswanadhan VN, Wendoloski JJ. A Knowledge-Based Approach in Designing Combinatorial or Medicinal Chemistry Libraries for Drug Discovery. 1. A Qual-itative and Quantitative Characterization of Known Drug Databases. Journal of Combinatorial Chemistry. 1999;1(1):55–68. PMID: 10746014. 10.1021/cc9800071.

53. Veber DF, Johnson SR, Cheng HY, Smith BR, Ward KW, Kopple KD. Molecular Properties That Influence the Oral Bioavailability of Drug Candidates. Journal of Medicinal Chemistry. 2002;45(12):2615–23. PMID: 12036371. 10.1021/jm020017n.

54. Egan WJ, Merz KM, Baldwin JJ. Prediction of Drug Absorption Using Multivariate Statistics. Journal of Medicinal Chemistry. 2000;43(21):3867–77. PMID: 11052792. 10.1021/jm000292e.

55. Muegge I, Heald SL, Brittelli D. Simple Selection Criteria for Drug-like Chemical Matter. Journal of Medicinal Chemistry. 2001;44(12):1841–6. PMID: 11384230. 10.1021/jm015507e.

56. Brenk R, et al. Lessons Learnt from Assembling Screening Libraries for Drug Discovery for Neglected Diseases. ChemMedChem. 2008;3(3):435–44. 10.1002/cmdc.200700139.

57. Baell JB, Holloway GA. New Substructure Filters for Removal of Pan Assay Interference Compounds (PAINS) from Screening Libraries and for Their Exclusion in Bioassays. Journal of Medicinal Chemistry. 2010 Apr;53(7):2719–40. 10.1021/jm901137j.

58. Romero, R., (2024). Tyrosine Kinases ligands with bioactivity data [Data set]. Kaggle. 10.34740/KAGGLE/DSV/8626441.

59. Romero, R. (2024). Bioactivity Deep Learning Classification Model. Zenodo. https://zenodo.org/doi/10.5281/zenodo.12682924.

60. Páll S, Zhmurov A, Bauer P, et al. Heterogeneous parallelization and acceleration of molecular dynamics simulations in GROMACS. J Chem Phys. 2020;153(13):134110. 10.1063/5.0018516.

61. Gerstner N, Kehl T, Lenhof K, et al. GeneTrail: A Framework for the Analysis of High-Throughput Profiles. Front Mol Biosci. 2021; 8:716544. 2021. 10.3389/fmolb.2021.716544.

62. Martens M, et al. WikiPathways: connecting communities. Nucleic Acids Research. 2020 11;49(D1):D613–21. 10.1093/nar/gkaa1024.

63. Kanehisa M, Furumichi M, Sato Y, Ishiguro-Watanabe M, Tanabe M. KEGG: integrating viruses and cellular organisms. Nucleic Acids Res. 2020 10;49(D1):D545–51. 10.1093/nar/gkaa970.

64. Milacic M. The Reactome Pathway Knowledgebase 2024. Nucleic Acids Research. 2024. 10.1093/nar/gkad1025.

65. Ashburner M, et al. Gene Ontology: tool for the unification of biology. Nature Genetics. 2000 May;25(1):25–9. 10.1038/75556.

66. Consortium TGO, Aleksander SA, Balhoff J, Carbon S, Cherry JM, Drabkin HJ, et al. The Gene Ontology knowledgebase in 2023. Genetics. 2023 03;224(1):iyad031. 10.1093/genetics/iyad031.

67. Lakkireddy Samyuktha, et al. In Silico Docking Studies of Vascular Endothelial Growth Factor-A (VEGFA): Possible Implications in Chronic Myeloid Leukemia (CML) Therapy, Current Proteomics 2021; 18 (4). 10.2174/1570164617999200929125324.

68. Meyer EA, Castellano RK, Diederich F. Interactions with aromatic rings in chemical and biological recognition. Angew Chem Int Ed Engl. 2003;42(11):1210–1250. 10.1002/anie.200390319

69. Lanzetta N, Giannozzi P, Crescenzi O, Improta R. π–π Interactions in Protein–Ligand Complexes: A Comprehensive Study Using Quantum Mechanical Methods. J Chem Inf Model. 2023;63(2):603–618. 10.1021/acs.jcim.2c01362.

70. Du QS, Long SY, Meng JZ, Huang RB. Energetic analysis of pi-pi stacking and cation-pi interactions in protein structures. Mol Phys. 2013;111(9-11):1550–1560. 10.1080/00268976.2012.758725.

71. Wheeler SE, Bloom JWG. Toward a more complete understanding of noncovalent interactions involving aromatic rings. J Phys Chem A. 2014;118(32):6133–6147. 10.1021/jp504415p.

72. Mohit Narwal, Jarkko Koivunen, et al, (2013) Discovery of Tankyrase Inhibiting Flavones with Increased Potency and Isoenzyme Selectivity Journal of Medicinal Chemistry 56 (20), 7880–7889 doi: 10.1021/jm401463y.

73. Vishnu Nayak Badavath, et al, (2015) Monoamine oxidase inhibitory activity of 2-aryl-4H-chromen-4-ones, Bioorganic Chemistry, Volume 58, Pages 72–80, ISSN 0045-2068, 10.1016/j.bioorg.2014.11.008.

74. Tamura H, et al. The systematic structure-activity relationship to predict how flavones bind to human androgen receptor for their antagonistic activity. Bioorg Med Chem. 2013 Jun 1;21(11):2968–74. doi: 10.1016/j.bmc.2013.03.060.

75. Muftuoglu Y, Mustata G. Pharmacophore modeling strategies for the development of novel nonsteroidal inhibitors of human aromatase (CYP19). Bioorg Med Chem Lett. 2010 May 15;20(10):3050–64. doi: 10.1016/j.bmcl.2010.03.113. Epub 2010 Apr 8. Erratum in: Bioorg Med Chem Lett. 2010 Aug 15;20(16):4955-9.

76. Stupp R, Mason WP, van den Bent MJ, et al. Radiotherapy plus concomitant and adjuvant temozolomide for glioblastoma. N Engl J Med. 2005;352(10):987–996. doi:10.1056/NEJMoa043330

77. Huang Y, Kim BYS, Chan CK, et al. Improving immune-vascular crosstalk for cancer immunotherapy. Nat Rev Immunol. 2018;18(3):195–203. doi:10.1038/nri.2017.145

78. Aldape K, Brindle KM, Chesler L, et al. Challenges to curing primary brain tumours. Nat Rev Clin Oncol. 2019;16(8):509–520. doi:10.1038/s41571-019-0177-5

79. Lu KV, Chang JP, Parachoniak CA, et al. VEGF inhibits tumor cell invasion and mesenchymal transition through a MET/VEGFR2 complex. Cancer Cell. 2012;22(1):21–35. doi:10.1016/j.ccr.2012.05.037

80. Wick W, Gorlia T, Bendszus M, et al. Lomustine and Bevacizumab in Progressive Glioblastoma. N Engl J Med. 2017;377(20):1954–1963. doi:10.1056/NEJMoa1707358

81. Hambardzumyan D, Gutmann DH, Kettenmann H. The role of microglia and macrophages in glioma maintenance and progression. Nat Neurosci. 2016;19(1):20–27. doi: 10.1038/nn.4185

82. Komohara Y, Ohnishi K, Kuratsu J, Takeya M. Possible involvement of the M2 anti-inflammatory macrophage phenotype in growth of human gliomas. J Pathol. 2008;216(1):15–24. doi: 10.1002/path.2370

83. Quail DF, Joyce JA. The microenvironmental landscape of brain tumors. Cancer Cell. 2017;31(3):326–341. doi: 10.1016/j.ccell.2017.02.009

84. Neftel C, Laffy J, Filbin MG, et al. An integrative model of cellular states, plasticity, and genetics for glioblastoma. Cell. 2019;178(4):835–849.e21. doi: 10.1016/j.cell.2019.06.024

85. Patel AP, Tirosh I, Trombetta JJ, et al. Single-cell RNA-seq highlights intratumoral heterogeneity in primary glioblastoma. Science. 2014;344(6190):1396–1401. doi: 10.1126/science.1254257

86. Deretic V. Autophagy in inflammation, infection, and immunometabolism. Immunity. 2021;54(3):437–453. doi: 10.1016/j.immuni.2021.01.018

87. Cavallaro U, Christofori G. Cell adhesion and signalling by cadherins and Ig-CAMs in cancer. Nat Rev Cancer. 2004;4(2):118–132. doi: 10.1038/nrc1276

88. Hambardzumyan D, Gutmann DH, Kettenmann H. The role of microglia and macrophages in glioma maintenance and progression. Nat Neurosci. 2016;19(1):20–27. doi: 10.1038/nn.4185

89. Quail DF, Joyce JA. The microenvironmental landscape of brain tumors. Cancer Cell. 2017;31(3):326–341. doi: 10.1016/j.ccell.2017.02.009

90. Paloneva J, Mandelin J, Kiialainen A, et al. DAP12/TREM2 deficiency results in impaired osteoclast differentiation and osteoporotic features. J Exp Med. 2003;198(4):669–675. doi: 10.1084/jem.20030827

91. Hu F, Dzaye O, Hahn A, et al. Glioma-derived versican promotes tumor expansion via glioma-associated microglial/macrophage Toll-like receptor 2 signaling. *Neuro Oncol*. 2015;17(2):200–210. doi: 10.1093/neuonc/nou137

92. Butowski NA. Immunotherapy for glioblastoma. Curr Neurol Neurosci Rep. 2017;17(4):29. doi: 10.1007/s11910-017-0738-z

93. Komohara Y, Ohnishi K, Kuratsu J, Takeya M. Possible involvement of the M2 anti-inflammatory macrophage phenotype in growth of human gliomas. J Pathol. 2008;216(1):15–24. doi: 10.1002/path.2370

94. Bergenfeldt M, Nielsen HJ, Kehlet H, et al. Acute-phase protein concentration and the systemic inflammatory response following surgery in patients with malignant disease. Br J Cancer. 2003;89(3):499–504. doi: 10.1038/sj.bjc.6601139

95. Brown NF, Carter TJ, Ottaviani D, Mulholland P. Harnessing the immune system in glioblastoma. Br J Cancer. 2018;119(10):1171–1181. doi: 10.1038/s41416-018-0265-8

