## Supplementary figures for "Identification of Potential Therapeutic Targets and Biomarkers for Glioblastomas Through Integrative Analysis of Gene Expression Data"

Supplementary figures for volcano plots and Venn diagrams of GEO studies, PPI networks ans enrichment analysis of downregulated hub genes.

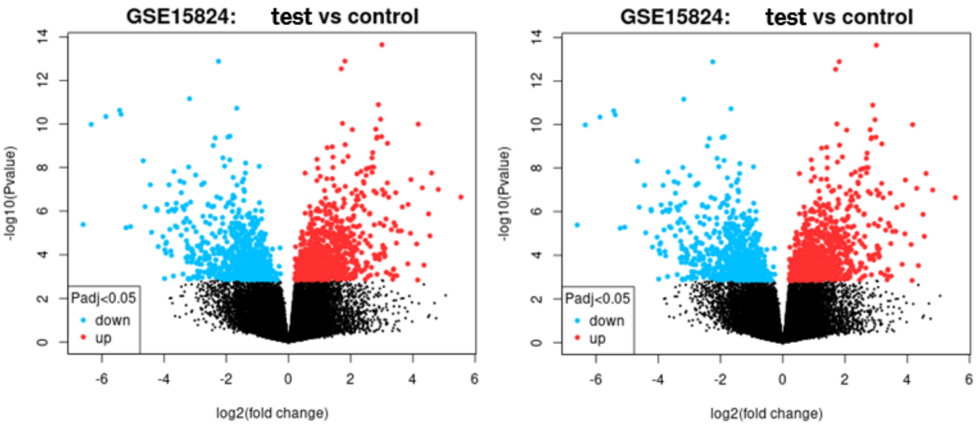


Figure 1. Volcano plot showing the dysregulated genes resulting from differential gene expression analysis, for both GEO studies employed.

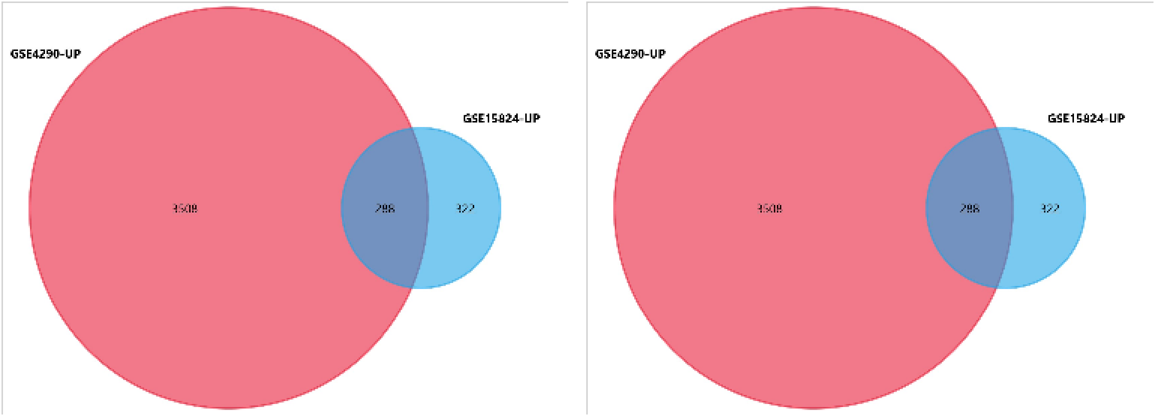


Figure 2. Venn Diagrams showing the number of common genes for up-regulated genes (left) and down-regulated ones (right) from GEO studies.

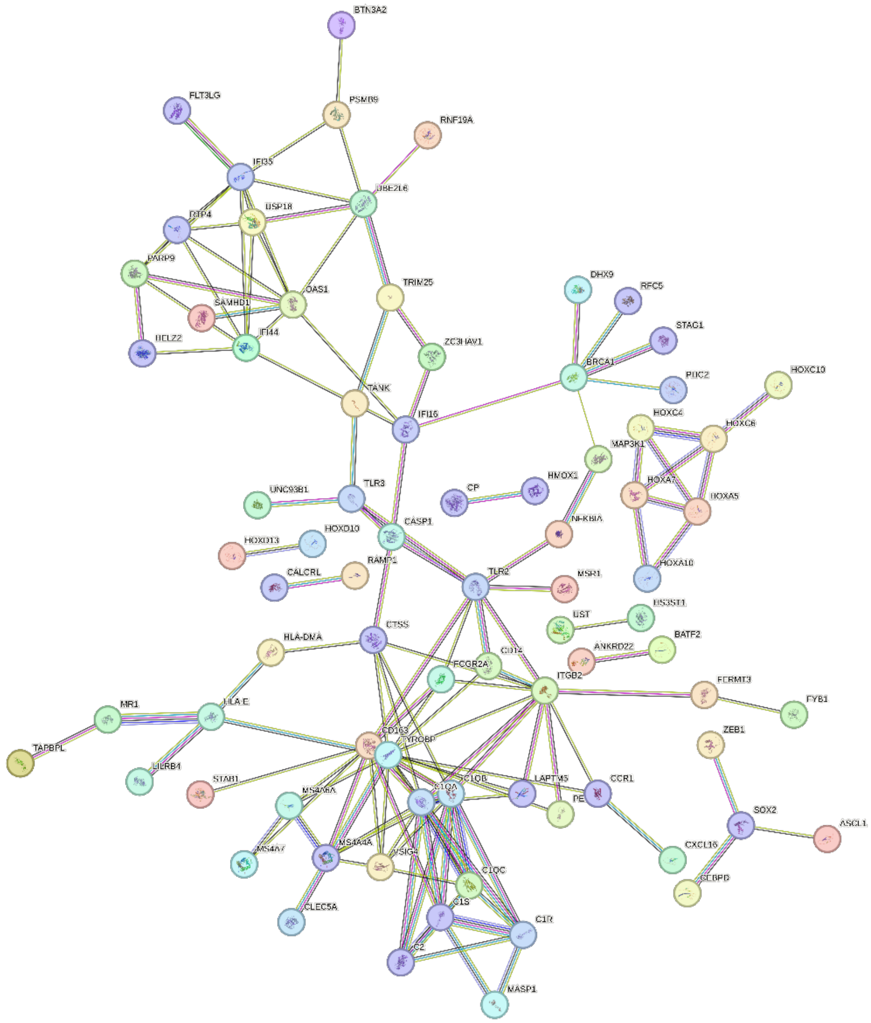

Figure 3. PPI network of upregulated genes from both databases. The interaction score was set to 0.7 (High). Number of nodes:172, number of edges: 35, average node degree: 1.57, avg. local clustering coefficient: 0.297, expected number of edges: 26, PPI enrichment p-value: < 1.0e-16.

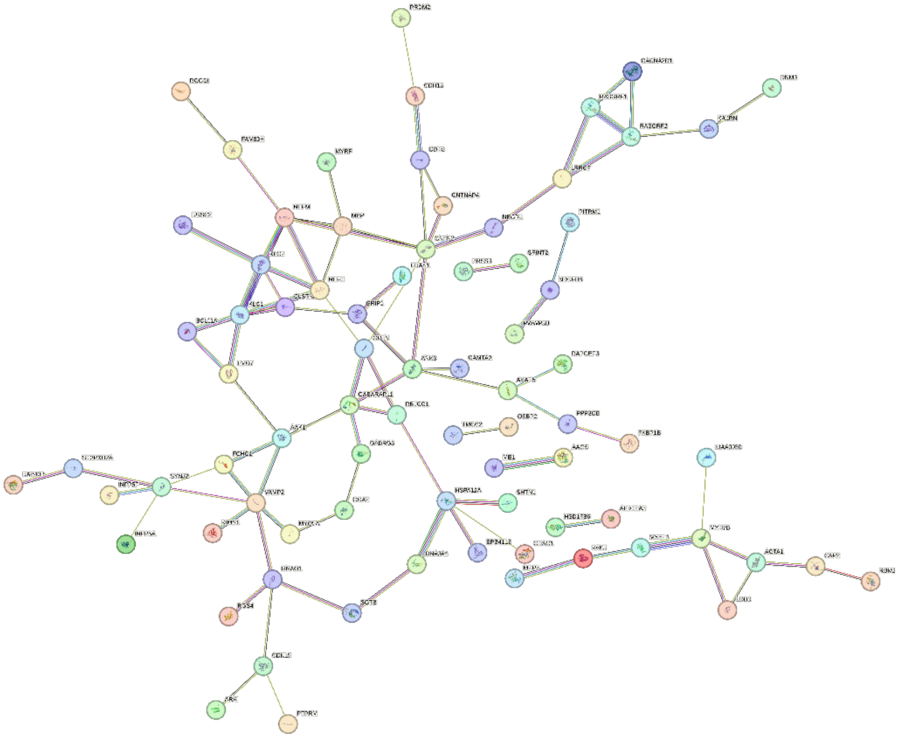

Figure 4. PPI network of upregulated genes from both databases. The interaction score was set to 0.4 (Normal). Number of nodes:147, number of edges: 91, average node degree: 1.24, avg. local clustering coefficient: 0.288, expected number of edges: 32, PPI enrichment p-value: < 1.0e-16.

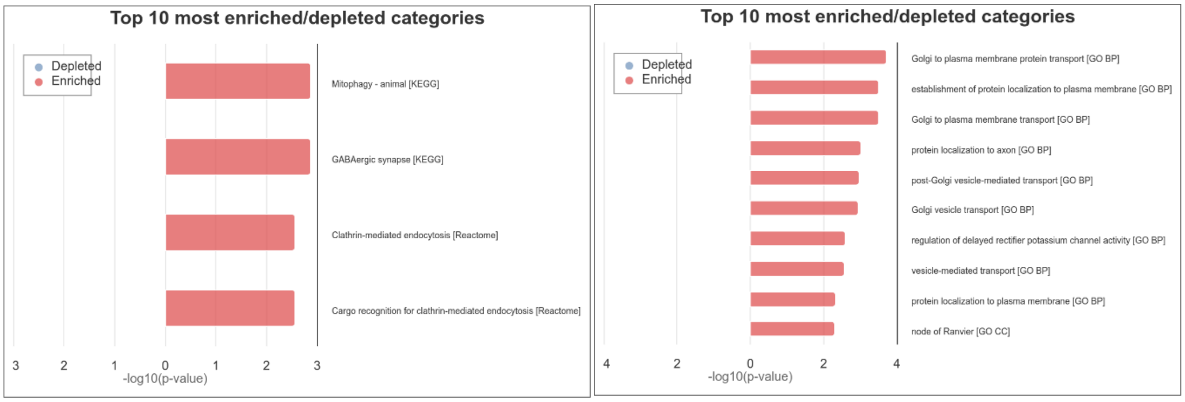


Figure 5. Top 10 enriched categories for pathways (left) and gene ontology (right) for downregulated genes in glioblastomas. The left panel illustrates the most enriched biological pathways related to down-regulated genes in glioblastomas, including "Mitophagy - animal," "GABAergic synapse," and "Clathrin-mediated endocytosis" (KEGG and Reactome databases), indicating a disruption in mitochondrial function, neurotransmission, and cellular transport mechanisms. The right panel shows the most enriched Gene Ontology (GO) categories, emphasizing cellular transport processes such as "Golgi to plasma membrane protein transport" and "establishment of protein localization to plasma membrane," along with processes related to neuronal functions and vesicle-mediated transport. These findings suggest that the down-regulation of these genes is associated with impaired cellular transport and synaptic function, which may contribute to the pathology of glioblastomas.
